# Kinetic Analysis of Lanthipeptide Cyclization by Substrate-Tolerant ProcM

**DOI:** 10.1101/2024.05.16.594612

**Authors:** Emily K. Desormeaux, Wilfred A. van der Donk

## Abstract

Lanthipeptides are ribosomally synthesized and post-translationally modified peptides characterized by the presence of thioether crosslinks. Class II lanthipeptide synthetases are bifunctional enzymes responsible for the multistep chemical modification of these natural products. ProcM is a class II lanthipeptide synthetase known for its remarkable substrate tolerance and ability to install diverse (methyl)lanthionine rings with high accuracy. Previous studies suggested that the final ring pattern of the lanthipeptide product may be determined by the substrate sequence rather than by ProcM, and that ProcM operates by a kinetically controlled mechanism, wherein the ring pattern is dictated by the relative rates of the individual cyclization reactions. This study utilizes kinetic assays to determine if rates of isolated modifications can predict the final ring pattern present in prochlorosins. Changes in the core substrate sequence resulted in changes to the reaction rates of ring formation as well as a change in the order of modifications. Additionally, individual chemical reaction rates were significantly impacted by the presence of other modifications on the peptide. These findings indicate that the rates of isolated modifications are capable of predicting the final ring pattern but are not necessarily a good predictor of the order of modification in WT ProcA3.3 and its variants.

## Introduction

Natural products are an abundant source of compounds with diverse bioactivities. Approximately two-thirds of all drugs approved between 1981 and 2019 are natural products or their derivatives.^1^ Ribosomally synthesized and post-translationally modified peptides (RiPPs) are a class of natural products that exhibit a wide array of biological activities.^2^ Biosynthetic gene clusters (BGCs) for these compounds contain a gene encoding a precursor peptide (LanA), that is post-translationally modified by biosynthetic enzymes to produce the mature natural product (Figure 1A).^3,4^ The precursor peptide is composed of an N-terminal leader region, which is typically important for enzyme recognition, and a C-terminal core region, which undergoes modification. After modification, the leader region is removed via proteases to yield the mature natural product.

**Figure 1.**
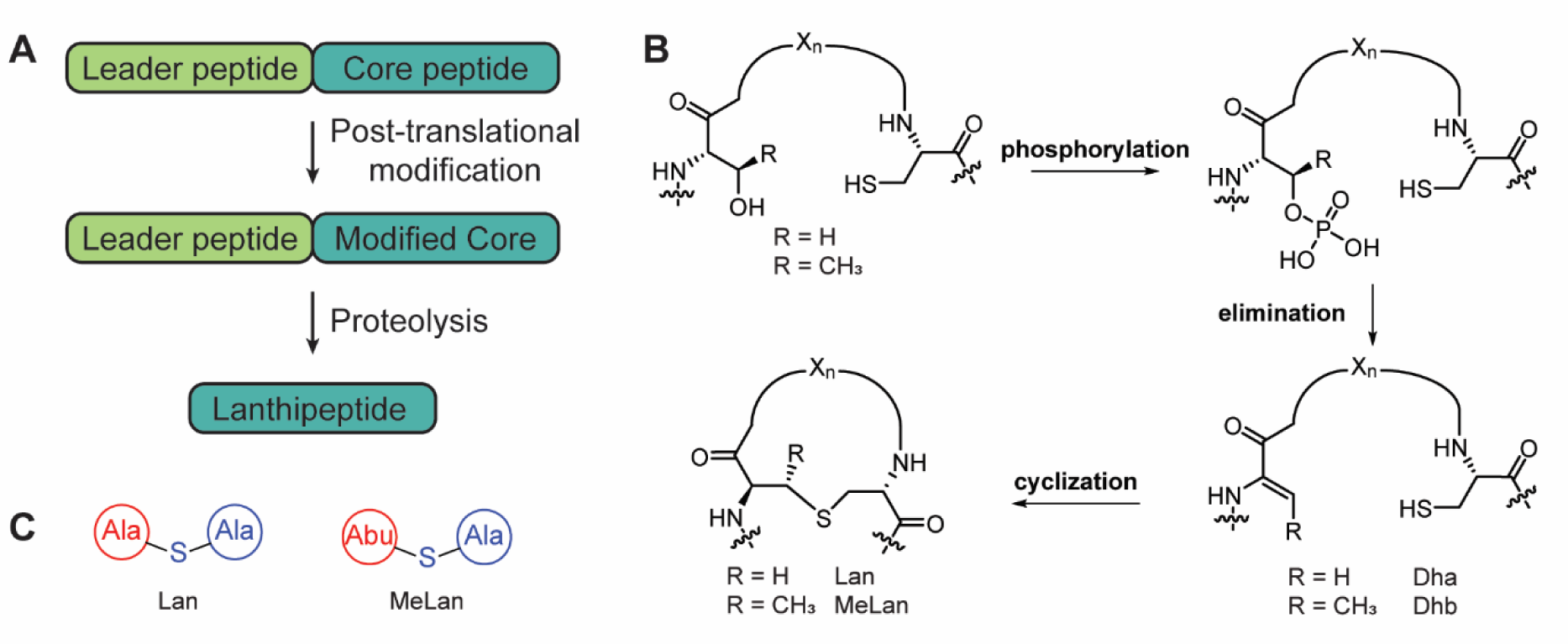
Biosynthetic pathways to lanthipeptides. **A)** Generic scheme for RiPP biosynthesis. **B)** Biosynthetic scheme for the post-translational modifications installed by class II lanthipeptide synthetases. **C)** Shorthand notation for lanthionine (Lan) and methyllanthionine (MeLan).

Lanthipeptides are a class of RiPPs characterized by the presence of lanthionine (Lan) and/or methyllanthionine (MeLan).^3^ These thioether cross-links are installed by (at least) five different lanthipeptide synthetase classes through the dehydration of Ser/Thr residues and addition of a Cys to the dehydrated residue.^4–10^ Class II lanthipeptide synthetases, LanMs, are single bifunctional enzymes that catalyze both the dehydration and the cyclization reactions (Figure 1B). The N-terminal dehydratase domain catalyzes the dehydration of Ser/Thr residues using ATP to first generate a phosphorylated peptide intermediate, which undergoes elimination in the same active site to yield dehydroalanine or dehydrobutyrine (Dha and Dhb, respectively).^11,12^ The C-terminal cyclization domain next catalyzes the Michael-type addition of a Cys thiol to the dehydrated residue to produce the (Me)Lan ring(s).^13^ Some class II lanthipeptide synthetases are found alongside a large number of precursor peptides (10-80)^14–17^ deviating from the majority of RiPP BGCs, which typically encode a single precursor gene. These gene clusters have been of particular interest for bioengineering efforts since the proteins have demonstrated relaxed substrate specificity.^17–25^

ProcM is a highly substrate-tolerant class II lanthipeptide synthetase discovered in the marine picocyanobacterium *Prochlorococcus* MIT9313.^14^ Along with the gene encoding ProcM, thirty putative genetically encoded precursor peptides, termed ProcAs, were identified within the genome.^14,18^ Of these precursors, eighteen ProcA peptides have been tested and were all modified by ProcM to produce polycyclic products with defined ring patterns termed prochlorosins (Pcns).^14,26,27^ Mechanistic studies performed on ProcM support a kinetically controlled mechanism, as the retro-Michael addition was not observed.^28,29^ Kinetic analysis performed on ProcM modification of the substrate ProcA2.8 support this hypothesis and suggests that the rates of each ProcM-catalyzed cyclization progressively decrease as the peptide acquires successive modifications.^30^

The ability of ProcM to produce Pcns with a wide diversity of ring patterns resulted in the utilization of ProcM for bioengineering efforts.^20–23^ Tested members from a ProcA2.8 analog library all formed the canonical ring pattern present in the wild-type (WT) peptide, whereas a library composed of ProcA3.3 variants produced a mixture of the native overlapping and non-native non-overlapping ring patterns (Figure 2).^21,23^ These findings support a model in which the final Pcn ring pattern is dictated by the precursor peptide sequence, which may be driven by conformational preferences.^31^ Cyclization of dehydrated ProcA3.3 also results in the alternative non-overlapping pattern in the absence of ProcM,^29^ similar to the predictions of computational models.^31^ These results suggest that while precursor peptide sequence is an important factor, that ProcM also plays some role in governing the final ring pattern.^32^

**Figure 2.**
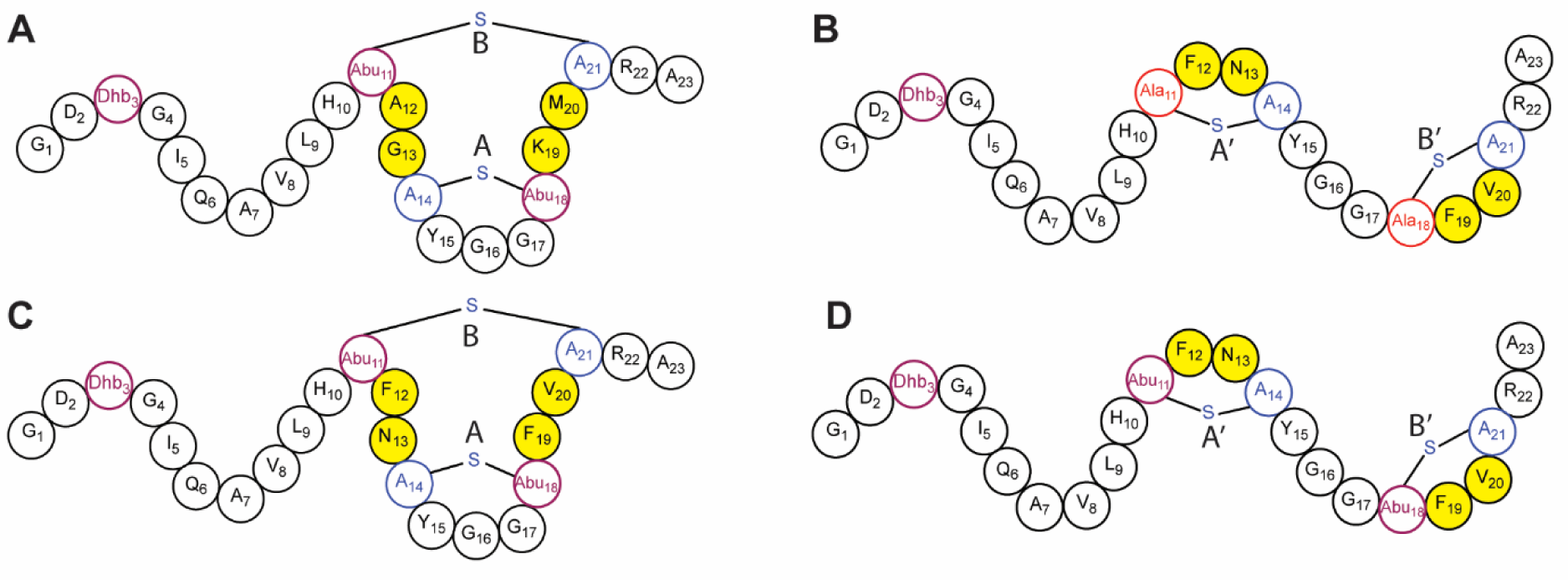
Structure of prochlorosin peptides represented by shorthand notation. **A)** Pcn3.3 WT with overlapping ring pattern containing A and B rings. **B)** Pcn3.3 variant Z with non-overlapping ring pattern containing A’ and B’ rings. **C/D)** MeLan-containing Pcn3.3 variant Z, which produces a mixture of the overlapping and non-overlapping ring patterns. The four positions that were randomized in the ProcA3.3 analog library are indicated in yellow.

To determine the importance of substrate sequence and ProcM influence on Pcn maturation, kinetic analysis was utilized in this work to characterize the maturation of WT ProcA3.3 (Figure 2A) as well as a ProcA3.3 variant (variant Z) that forms a non-overlapping ring pattern (Figure 2B).

In addition to these analyses, we measured cyclization kinetics for individual rings to determine if the isolated cyclization rates can be used to predict the final ring pattern. We found that the mutation of Thr to Ser and vice versa has a large impact on both the dehydration and cyclization steps during peptide modification. Rates of formation of individual rings could be used to predict the final ring pattern in the case of WT ProcA3.3 and variant Z.

## Methods

### ProcM Kinetic Assays

ProcA peptides and ProcM were expressed and purified as previously described.^30,31^ ProcM (2 or 4 µM) and ProcA peptide (80, 120, or 160 µM) were preincubated separately at 25 °C for 1 h in 750 µL of reaction mixture (5 mM ATP, 0.17 mM ADP, 5 mM MgCl_2_, 100 mM HEPES, 0.1 mM TCEP, pH 7.5) to allow for equilibration prior to initiation of the reaction. The reaction mixture was altered from previous methods with the addition of ADP.^30^ ADP was added to the reaction mixture as it helps to limit build-up of phosphorylated intermediates and allows intermediates to rebind ProcM when present; these intermediates, when observed in previous publications, often act as dead-end products in the absence of ADP that are not converted to the dehydrated intermediate.^30^ Under physiological conditions ADP would be present in the ATP/ADP ratio used here.^33^ The peptide and protein samples were mixed thoroughly to initiate a reaction. At desired time points (2, 4, 6, 8, 10, 15, 18, 20, 30, 45, 60, 75, 90, 120, 180, and 240 min), 80 µL aliquots were removed from the reaction. The reaction aliquot was quenched using 900 µL of quench buffer (111 mM citrate, 1.11 mM EDTA, pH 3.0) and placed on ice until completion of all time points. Quenched samples were reduced with the addition of 100 µL of 100 mM TCEP, and the samples were incubated at 25 °C for 10 min. The pH was then adjusted to approximately 6.3 with the addition of 40-45 µL of 5 M NaOH. Non-cyclized cysteine residues were then alkylated by the addition of 11 µL of 1 M *N*-ethylmaleimide (NEM) in EtOH, and the alkylation reaction was allowed to proceed for 10 min at 37 °C before quenching with the addition of 11 µL of formic acid. The solvent for each sample was then exchanged to water via centrifugal filtration using Amicon 3 kDa filters. The centrifugation step was altered from published protocols due to the discontinuation of Vydac BioSelect columns.^30^ Samples were lyophilized and stored at -80°C until analysis.

### Liquid Chromatography Mass Spectrometry-Kinetic samples

Dried samples were resuspended in 50 µL of water and the total peptide concentration was estimated via UV-vis absorption at 280 nm. Samples were then diluted to 15 µM total peptide in LCMS solvent (50% ACN in H_2_O, 0.1% formic acid). Samples (10 µL) were injected into an Acquity UPLC BEH C8 1.7 µm column (1.0 x 100 mm) coupled to a Waters quadrupole/ time-of-flight (Q/ToF) Synapt-G1 series mass spectrometer. The samples were analyzed with a flow rate of 0.4 mL/min and the peptides were eluted using the following protocol: 3% B held for 2 min, 3-97 %B over 3 min, and held at 97% B for 3 min followed by re-equilibration of the column to starting conditions (A: 0.1% formic acid in water, B: 100% ACN, 0.1% formic acid). All peptides eluted between 5 and 6 min in a single broad peak. All samples were run in a randomized order and 10 µL injections of water were run between each sample to prevent cross contamination. The mass spectrometry conditions were as follows: positive ion mode, V optics, capillary voltage = 3.0 kV, cone gas = 180 L/h, desolvation gas = 600 L/h, source temperature = 120 °C, desolvation temperature = 200 °C. The detector was set to a *m/z* window of 800-1800 Da with a 1 s scan time. External calibration was performed using a 0.1% phosphoric acid standard. Identity of each ion was determined by HRMS (Table S1) and MS/MS fragmentation patterns.

The total ion chromatogram (TIC) for each sample was used to determine *m/z* values for ions of interest. These m/z values were then used along with a mass window around the center of the most intense isotopic peak to generate extracted ion chromatograms (EICs). Integration of the EICs using Waters MassLynx software was performed, and the peak areas were then used to calculate the fractional abundance of each peptide species using previously published methods.^30^

### Simulation of Kinetic Data

The calculated time-dependent fractional abundances for each sample were imported into KinTek Explorer,^34,35^ and the standard deviation for each ion during the time course was estimated via fitting to exponential equations using the “aFit” module. A kinetic model was then designed in KinTek and the time courses were used to fit the model to the collected data. Simplifying assumptions were used to create a minimal kinetic model that described the data collected, following previously published methods.^30^ Binding constants of ProcA peptides to ProcM from previous studies were used to approximate binding for all peptides utilized in this study. Goodness-of-fit of the models was evaluated within the KinTek software by χ^2^/DoF (the degrees of freedom), wherein for a good fit, χ^2^/DoF approaches 1. DoF is defined as n-1 where n is the number of variable parameters in the designed kinetic model.

The FitSpace Explorer module of KinTek was then utilized to generate confidence contours and evaluate the quality of data fitting. A confidence contour is a plot of the χ^2^ / χ^2^ value as a function of a variable parameter and can provide information on how well constrained that parameter is by the given model as well as providing well-defined upper and lower confidence intervals. This type of analysis is helpful when the distribution of errors is uneven (i.e., well-defined lower boundary and poorly defined upper boundary). The parameter boundaries for the rates in the kinetic models for WT ProcA3.3 and ProcA3.3 variant Z and their single ring analogs were calculated using χ^2^ thresholds of 0.83 and 0.96 respectively, which were the recommended boundaries by FitSpace Explorer.

### Liquid Chromatography Mass Spectrometry – Fragmentation analysis

To determine the location of modifications in partially modified ProcA3.3 variant Z, the remaining sample after kinetic analysis of the 10-minute time point of the 80 µM kinetic reaction was analyzed using previously published methods.^31^ To determine the final ring pattern of fully modified ProcA3.3 variants, the remaining sample after kinetic analysis of the end point of the 40 µM kinetic reaction was analyzed using previously published methods.^31^

## Results and Discussion

### ProcA3.3 WT Modification by ProcM

The maturation of ProcA3.3 WT (Figure 2A) involves the dehydration of three residues, Thr3, Thr11, and Thr18 as well as the cyclization of two overlapping MeLan rings, the A ring (formed between Cys14 and Thr18) and B ring (formed between Thr11 and Cys21). A previous study showed that the A ring forms before the B ring,^28^ but the timing of dehydration compared to cyclization was not determined. For the current kinetic analysis, the reaction mixture was subjected to NEM alkylation of unreacted Cys residues at various time points, resulting in uncyclized intermediates that are differentiated from cyclized peptides via a mass shift. Relevant peptides were then monitored via LC-MS analysis and fractional abundances were used to monitor the appearance and consumption of peptides (Figure 3).^30^

**Figure 3.**
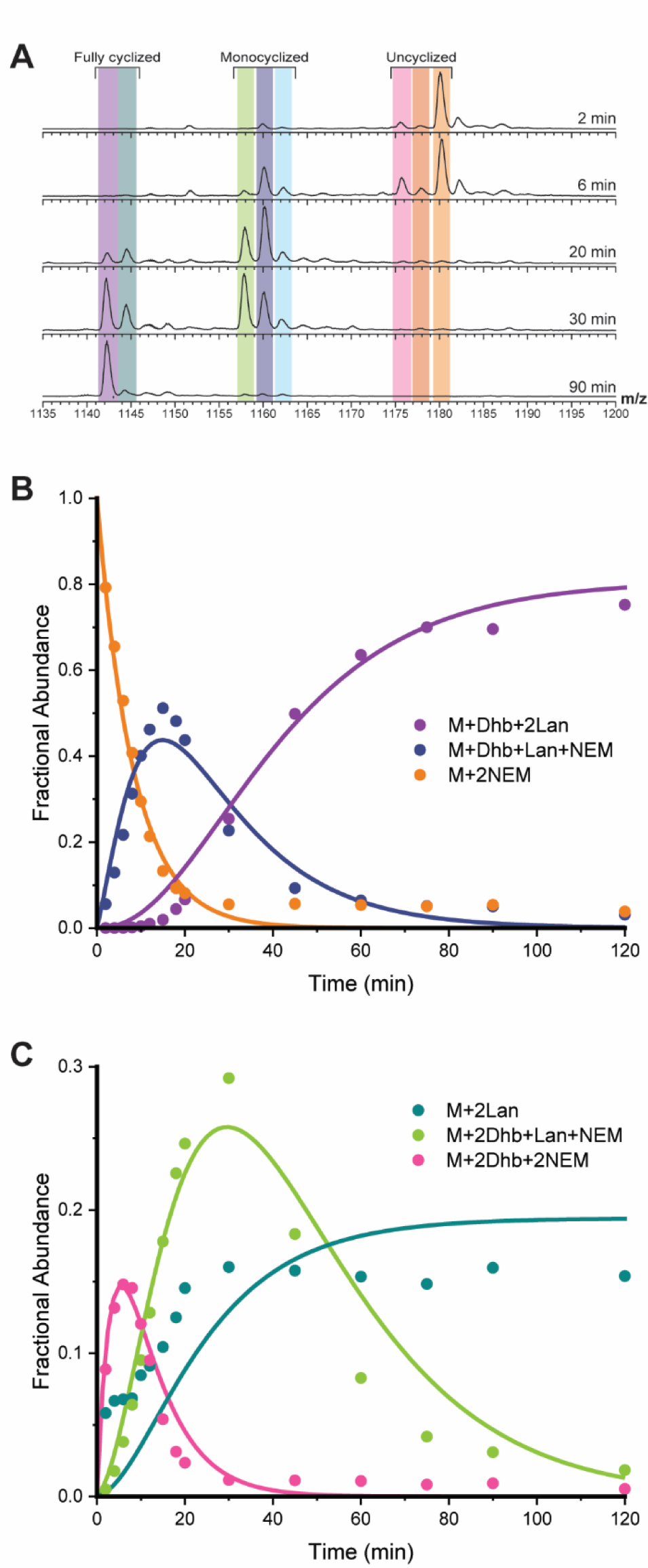
ProcA3.3 WT kinetic analysis. **A)** Representative ESI mass spectra of the 8+ charge state of ProcA3.3 WT at multiple time points. For each trace, the y-axis is scaled to the intensity of the highest peak present. The sample was treated with NEM to interrogate the cyclization state of the Cys residues. Coloring of each ion: light orange, unmodified peptide (M + 2 NEM); dark orange, once dehydrated peptide (M + Dhb + 2 NEM); pink, twice dehydrated peptide (M + 2 Dhb + 2 NEM); light blue, once dehydrated and cyclized peptide (M + MeLan + 1 NEM); dark blue, twice dehydrated and once cyclized peptide (M + Dhb + MeLan + 1 NEM); green, thrice dehydrated and once cyclized peptide (M + 2 Dhb + MeLan + 1 NEM); teal, twice dehydrated and twice cyclized peptide (M + 2 MeLan); purple thrice-dehydrated and twice cyclized peptide (M + Dhb + MeLan). **B)** Time course showing unmodified peptide (orange), peptide containing one MeLan ring and one additional dehydration (dark blue), and the final product (purple; M – 3 H2O). **C)** Time course for minor observed intermediates including non-cyclized peptide with two dehydrations (pink), peptide containing one MeLan ring with two dehydrations (green), and peptide containing two MeLan rings with no additional dehydrations (teal). Reaction for all panels were performed with 60 µM ProcA3.3 WT and 2 µM ProcM.

The reaction with WT ProcA3.3 was complete after approximately 45 min when only three-fold dehydrated, and twice cyclized peptide (purple) and two-fold dehydrated, and twice cyclized peptide (teal) were detected (Figure 3A). The main intermediates observed are two- and three-fold dehydrated peptide that have a single ring formed (dark blue and light green), and twice dehydrated peptide with no rings (pink). Intermediates containing a single dehydration (dark orange) and peptide containing a single methyllanthionine (light blue) were observed in low abundance. These intermediates were not used for the final minimal kinetic model. The peptide intermediate containing two MeLan rings and lacking the third dehydration (teal) was modeled as an end product as buildup of this intermediate remained through the completion of the reaction (Figure 3C, teal). We hypothesize that the twice cyclized product is unable to reach the dehydration active site of ProcM.

A minimal kinetic model representing the processing of WT ProcA3.3 by ProcM is depicted in Scheme 1A. Rate constants were determined for each chemical step via simulation of the data in Figures 3 and S1 using KinTek Explorer as described in the methods section. Several simplifying assumptions were used for the creation of this minimal model as described previously.^30^ Binding constants obtained in a previous study for one of the substrates were used for all peptides and intermediates. This assumption is based on the high sequence conservation of the leader region for ProcA substrates, and the major contribution of the leader peptide to the binding affinity for ProcM. Additionally, all chemical steps are assumed and modeled as irreversible. These assumptions are supported by dehydration being driven by ATP hydrolysis and ProcM having been demonstrated not to catalyze retro-Michael additions.^29^ To increase confidence the model was used for a global fit for multiple experiments conducted with varying concentrations of substrate and enzyme (Figure S1). The binding and chemical rate constants are provided in Table 1.

**Scheme 1.**
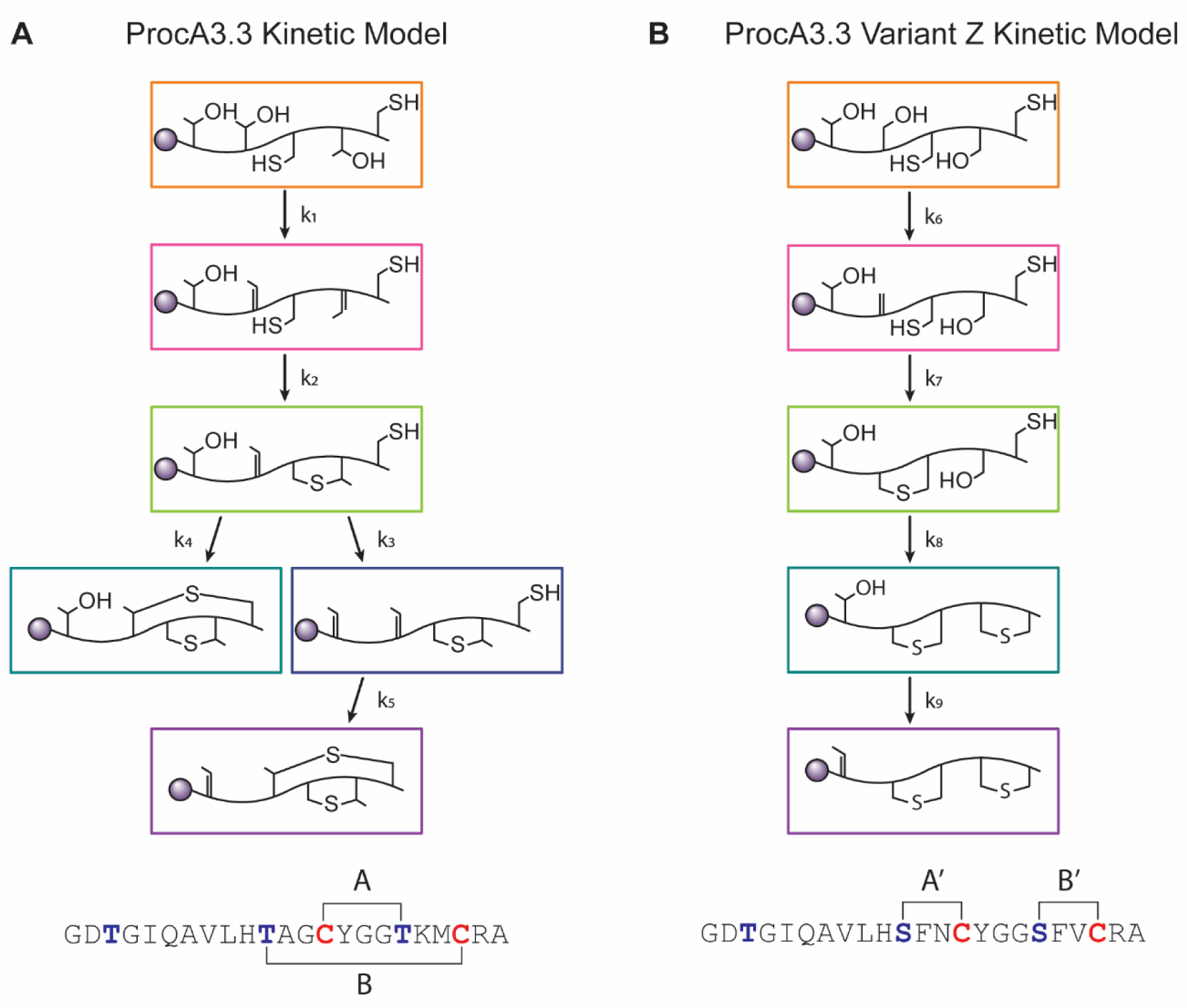
Kinetic models for WT ProcA3.3 and variant Z. A) Proposed minimal kinetic model for ProcA3.3 WT. Reactions for the initial two dehydrations were combined because the singly dehydrated intermediate is not observed in significant quantities under the reaction conditions. **B)** Proposed minimal kinetic model for ProcA3.3 variant Z. Notable differences are installation of a lanthionine ring before the second and third dehydrations occur. The sequences for both peptides are shown below their respective kinetic model.

**Table 1.**
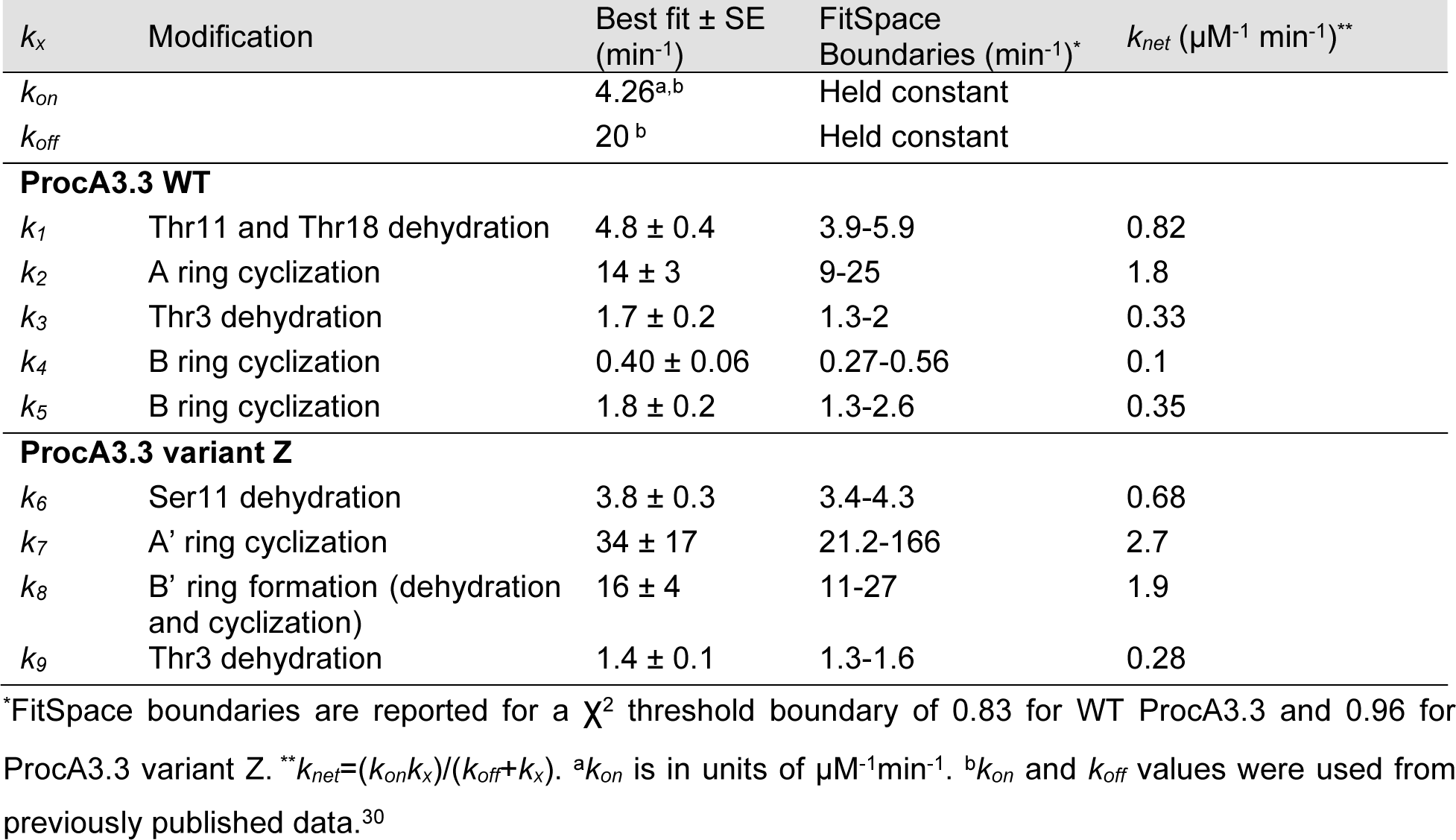
Summary of Simulated Rate Constants for Fully Modified Substrates.

| $k_x$ | Modification | Best fit ± SE (min<sup>-1</sup>) | FitSpace Boundaries (min<sup>-1</sup>)<sup>*</sup> | $k_{net}$ (μM<sup>-1</sup> min<sup>-1</sup>)<sup>**</sup> |
| --- | --- | --- | --- | --- |
| $k_{on}$ | | 4.26<sup>a,b</sup> | Held constant | |
| $k_{off}$ | | 20<sup>b</sup> | Held constant | |
| **ProcA3.3 WT** |  |  |  |  |
| $k_1$ | Thr11 and Thr18 dehydration | 4.8 ± 0.4 | 3.9-5.9 | 0.82 |
| $k_2$ | A ring cyclization | 14 ± 3 | 9-25 | 1.8 |
| $k_3$ | Thr3 dehydration | 1.7 ± 0.2 | 1.3-2 | 0.33 |
| $k_4$ | B ring cyclization | 0.40 ± 0.06 | 0.27-0.56 | 0.1 |
| $k_5$ | B ring cyclization | 1.8 ± 0.2 | 1.3-2.6 | 0.35 |
| **ProcA3.3 variant Z** |  |  |  |  |
| $k_6$ | Ser11 dehydration | 3.8 ± 0.3 | 3.4-4.3 | 0.68 |
| $k_7$ | A' ring cyclization | 34 ± 17 | 21.2-166 | 2.7 |
| $k_8$ | B' ring formation (dehydration and cyclization) | 16 ± 4 | 11-27 | 1.9 |
| $k_9$ | Thr3 dehydration | 1.4 ± 0.1 | 1.3-1.6 | 0.28 |

<sup>\*</sup>FitSpace boundaries are reported for a $\chi^2$ threshold boundary of 0.83 for WT ProcA3.3 and 0.96 for ProcA3.3 variant Z. <sup>\*\*</sup> $k_{net}=(k_{on}k_x)/(k_{off}+k_x)$ . <sup>a</sup> $k_{on}$ is in units of μM<sup>-1</sup>min<sup>-1</sup>. <sup>b</sup> $k_{on}$ and $k_{off}$ values were used from previously published data.<sup>30</sup>

The rate constants associated with cyclization, *k_2_* for ring A and *k_4/5_* for ring B, follow a similar trend to that of ProcA2.8,^30^ in which the first cyclization reaction is considerably faster than that of the second ring. The second cyclization reaction is modeled to occur from two possible intermediates; cyclization from a twice-dehydrated intermediate (*k_4_*) appears to produce a product that cannot be dehydrated a third time whereas cyclization from a three-times dehydrated peptide (*k_5_*) produces the fully modified product. The rate constants for these reactions were different (*k_4_* = 0.40 min^-1^, and *k_5_* = 1.8 min^-1^).

### ProcA3.3 Variant Z Modification by ProcM

The ProcA3.3 variant (Figure 2) used in this study was selected from a library of ProcA3.3 analogs made to disrupt protein-protein interactions.^21,23^ Library members contained two variations compared to WT. Firstly, the analogs featured T11S and T18S mutations that result in the formation of two Lan rings rather than MeLan rings. Secondly, the members of the library were randomized at four positions (shown in yellow in Figure 2) to assess the effect on cyclization pattern. The library member chosen for kinetic analysis in this study (termed variant Z) was fully modified by ProcM and produced almost exclusively the non-overlapping ring pattern. Under the reaction conditions provided in the methods section, only intermediates corresponding to the non-overlapping ring pattern were observed by tandem-MS analysis (Figure S2). Therefore, variant Z was a good candidate for kinetic analysis to understand the origins of the different cyclization site selectivity.

To generate a minimal kinetic model for ProcA3.3 variant Z, intermediates were again monitored by LC-MS analysis (Figure 4), and tandem MS/MS analysis was utilized to determine the location of the modifications (e.g. Figure S2). Tandem MS/MS analysis with the singly cyclized intermediate present at the 10-minute time point of a reaction containing 80 µM ProcA3.3 variant Z and 2 µM ProcM demonstrated that the NEM adduct was present on Cys21 and that dehydration occurred on Ser11, indicative of A’ ring formation (Figure S2). No ions were observed to suggest the presence of dehydration at other positions nor were NEM adducts detected at Cys14.

**Figure 4.**
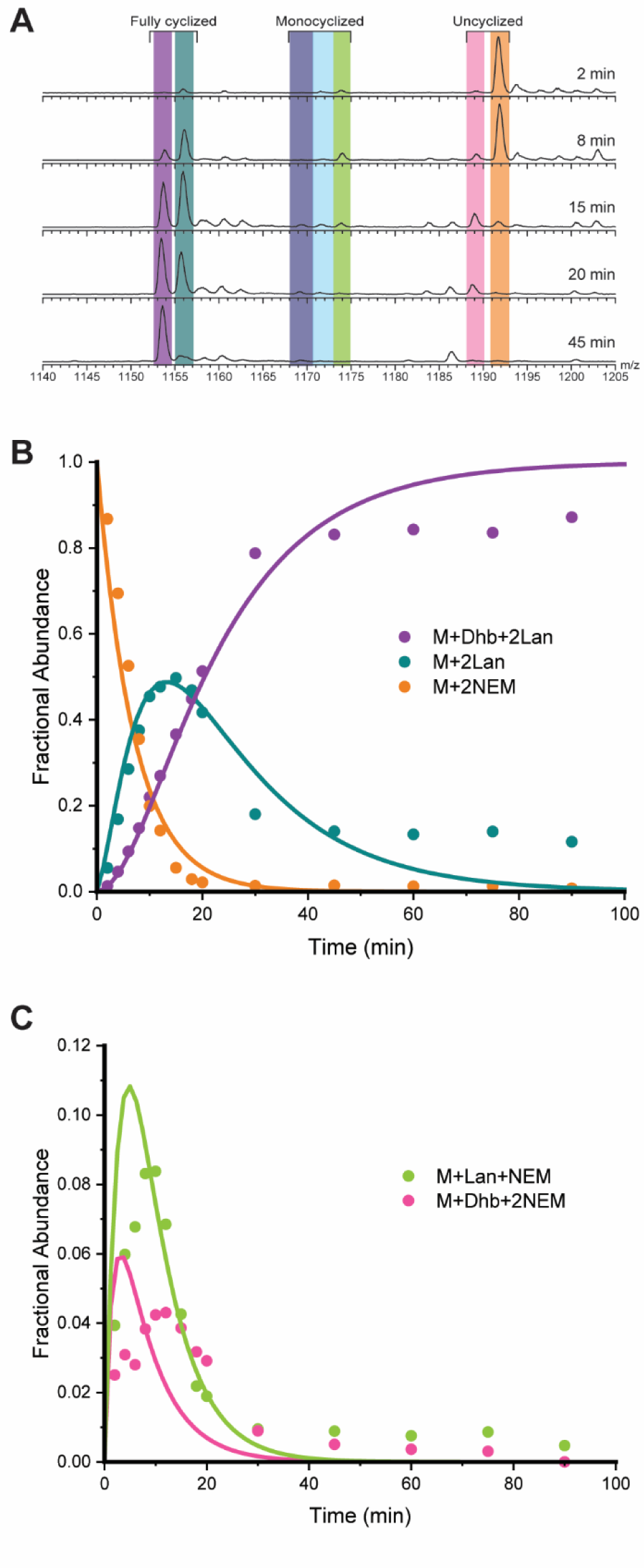
ProcA3.3 Variant Z Kinetic Analysis. **A)** Representative ESI mass spectra of the 8+ charge state of ProcA3.3 variant Z at multiple time points. Time course for the major species of the ProcA3.3 variant kinetic reaction with fractional abundances. Coloring of each ion: light orange, unmodified peptide (M + 2 NEM); pink, once dehydrated peptide (M + Dha + 2 NEM); green, once dehydrated and once cyclized peptide (M + Lan + 1 NEM); light blue, twice dehydrated and cyclized peptide (M + Dha + Lan + 1 NEM); dark blue, thrice dehydrated and once cyclized peptide (M + 2 Dhx + Lan + 1 NEM); teal, twice dehydrated and twice cyclized peptide (M + 2 Lan + 1 NEM); purple thrice-dehydrated and twice cyclized peptide (M + Dhb + 2 Lan). **B)** Time course for species including unmodified peptide (orange), peptide containing two Lan rings (cyan), and the final product (purple). **C)** Time course for minor species including non-cyclized peptide with one dehydration (pink) and peptide containing one Lan ring (green). Reaction conditions for all panels were performed with 40 µM ProcA3.3 variant Z and 2 µM ProcM.

Reactions were monitored, and fractional abundances were calculated for all relevant species. Two intermediates, one containing a single Lan ring and a single additional dehydration (light blue) and one containing a single Lan ring and two additional dehydrations (dark blue) were observed in low abundance. These species accounted for only approximately four percent of the total peptide concentration at their highest abundance. These intermediates were not used in the final minimal kinetic model. The simplifying assumptions described above were again used to develop a model for variant Z. Rate constants were determined for each chemical step shown in Scheme 1B via simulation of data shown in Figure 4 and S3 using KinTek Explorer as described in the methods section. The binding and chemical rate constants are present in Table 1.

The minimal kinetic model for modification of ProcA3.3 variant Z is depicted in Scheme 1B. The formation of the A’ ring indicates that the substitutions in the WT precursor sequence resulted in several changes in the order of modification of the peptide by ProcM. Most importantly, the A’ ring is formed prior to the second dehydration in this variant, which is unique compared to all previously characterized natural ProcA peptides including ProcA3.3, ProcA2.8, and ProcA1.1.^28,30,31^ This formation of the A’ ring sets the substrate on course to form a non-overlapping ring pattern.

For fully modified peptide, the dehydration rates of ProcA3.3 variant Z are comparable to those of ProcA3.3 WT. The first step in the kinetic model for each peptide, *k_1_* representing the dehydration of both Thr18 and Thr11 in WT, and *k_6_* representing the dehydration of Ser11 in variant Z, have rates of 4.8 and 3.8 min^-1^, respectively. The final dehydration step of Thr3, *k_3_* for ProcA3.3 WT and *k_9_* for ProcA3.3 variant Z, were 1.7 and 1.4 min^-1^, respectively. The dehydration steps having comparable rate constants suggests that the presence or lack of the methyl group on the Ser vs Thr does not alter the rate of chemical modification for these substrates.

The rate constants associated with cyclization for ProcA3.3 variant Z, *k_7_* for the A’ ring and *k_8_* for the B’ ring (34 and 16 min^-1^ respectively), follow a similar trend to that of other characterized lanthipeptides, including WT ProcA3.3 and WT ProcA2.8,^30^ wherein the first ring formation for variant Z has a larger rate constant compared to the second. The rate of B’ ring formation is a coupled rate constant of Ser18 dehydration and B’ ring formation. The absence of a dehydrated intermediate indicates that the cyclization rate is greater than dehydration, with the *k_8_* fit value limited by the rate of dehydration. It is therefore possible that the cyclization rate of the B’ ring is greater than that of the A’ ring. As noted in the model, the main finding of these kinetic studies is that the fast cyclization of the A’ ring even before dehydration of Ser18 precludes A-ring formation and leads to a non-overlapping ring pattern.

### Dhb-forming ProcA3.3 Variant Z Modification by ProcM

Variant Z has two types of changes compared to ProcA3.3. In order to independently determine the effect of the T11S and T18S mutations compared to the amino acid substitutions within the rings (A12F, G13N, K19F, and M20V) on the final ring patterns, a ProcA3.3 variant Z analog was generated that harbors S11T and S18T mutations (Figure 2C,D). This analog leads to methyllanthionines similar to the WT ProcA3.3 and the changes compared to WT are just the substitutions within the ring. This substrate was utilized for *in vitro* reactions with ProcM under the same conditions as WT and variant Z and subjected to LC-MS and tandem MS analysis as described in the methods section. The total ion chromatographs (TIC) of the product peptide indicated two distinct peaks that correspond to overlapping and non-overlapping ring patterns based on MS/MS fragmentation analysis (Figure S4). Several of the intermediate species were also subjected to MS/MS in an attempt to determine the location of modifications. The analysis indicated a mixture of peptide modification sites. As a result, the analog of ProcA3.3 variant Z was not subjected to kinetic analysis, as it would be impossible to define a minimal kinetic model without knowing distinct modification sites. Similarly, we also generated another analog of variant Z that retained the Ser residues at positions 11 and 18, but that had all other residues the same as WT ProcA3.3. Once again, this analog resulted in a product mixture of overlapping and non-overlapping rings (Figure S5) that was not further investigated. Collectively, these results show that the change from overlapping to non-overlapping ring patterns of the final products in going from WT to variant Z is caused by both the switch from MeLan to Lan rings, and the substitution of residues in the rings.

### Single ring cyclization kinetics

The ring pattern of the product(s) formed by ProcM from ProcA3.3 and its variants is determined by the relative rates of formation of the A, A’ and B’ rings. If the A ring is formed faster than both the A’ and B’ rings, the overlapping ring pattern will be formed. Conversely, if either the A’ or the B’ ring is formed faster than the A ring, the non-overlapping ring pattern will be generated. To determine if patterns for fully modified substrate peptides could be predicted by the relative rates of ring formation in isolation, peptides were designed to report on only one of the four possible rings per peptide (Figure S6). In these variants, all modifiable residues not involved in the ring of interest were mutated to alanine. We elected to do this only for the WT and variant Z ProcA3.3 peptides since they form single products. The variants were subjected to the same reaction conditions with ProcM and the reactions were analyzed via LC-MS.

### Single ring cyclization kinetics for ProcA3.3 WT

Single ring cyclization kinetic data were collected for mutants of the WT ProcA3.3 scaffold that were only able to create the A, B, A’ and B’ rings (Figure S6). Peaks corresponding to unmodified, dehydrated, and cyclized peptides were monitored and fractional abundance was again calculated to monitor the reaction progression (Figure 5 and Figures S7-S9). These data were used to generate a minimal kinetic model in which precursor peptide is modified to a MeLan ring containing peptide through a dehydration step, *k_dehy_*, and a cyclization step, *k_cycl_*(Scheme 2A). The use of previously determined binding constants was again applied to these models.

**Figure 5.**
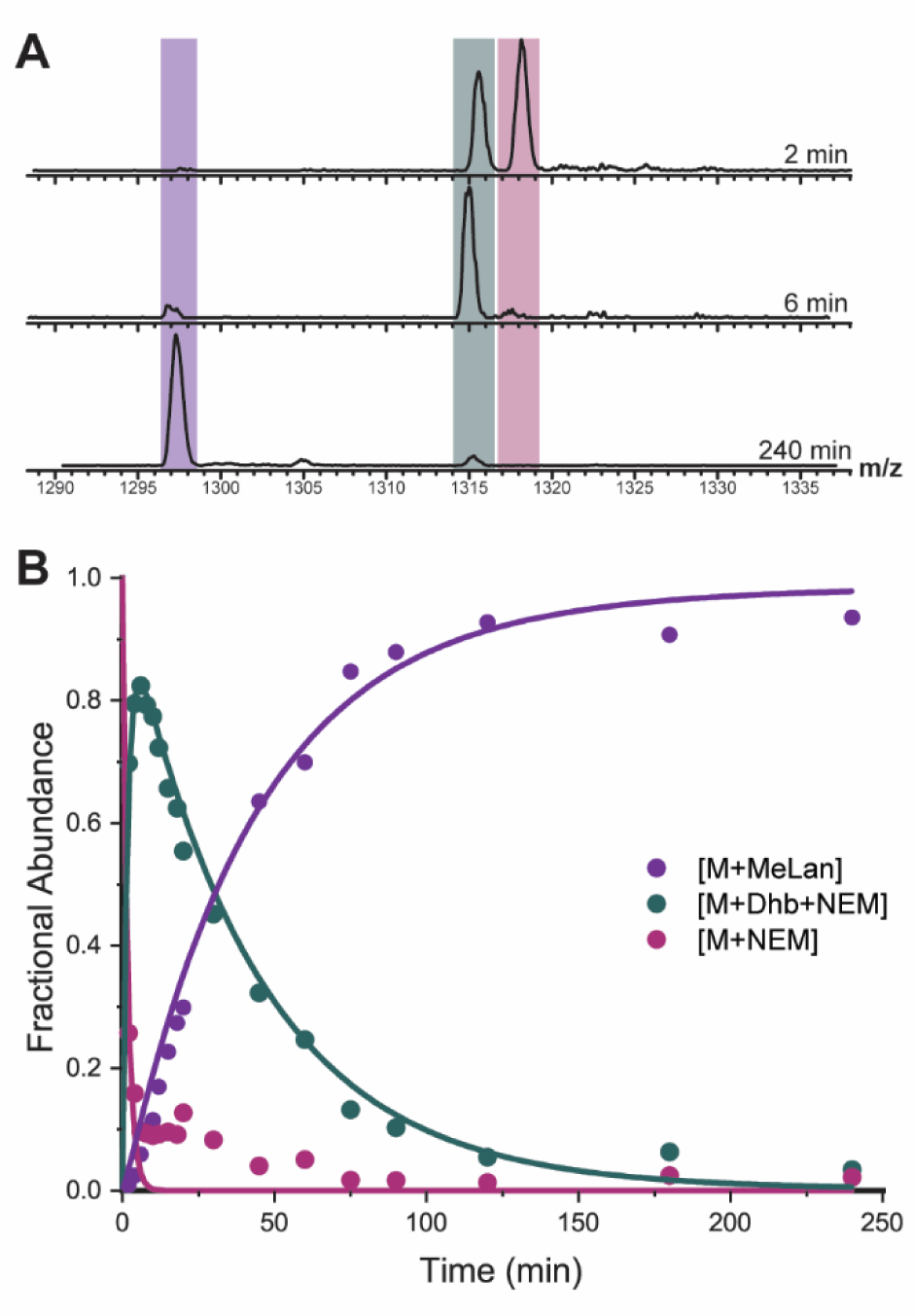
Representative example of kinetic data and fit for a ProcA3.3 variant that makes a single ring (B ring). Reactions for this figure were performed with ProcA3.3 B ring (ProcA3.3 T3A/C14A/T18A) at 40 µM substrate and 2 µM ProcM concentrations. **A)** Representative ESI mass spectra of the 7+ charge state of the variant that forms the ProcA3.3 B ring at multiple time points. Coloring of each ion: pink, unmodified peptide (M + NEM); teal, dehydrated peptide (M + 1 Dhb + NEM); purple, cyclized peptide (M + MeLan). **B)** Time course for the intermediates and product of the ProcA3.3 B ring formation with fractional abundances for unmodified peptide (pink), dehydrated peptide (teal), and cyclized peptide (purple).

**Scheme 2.**
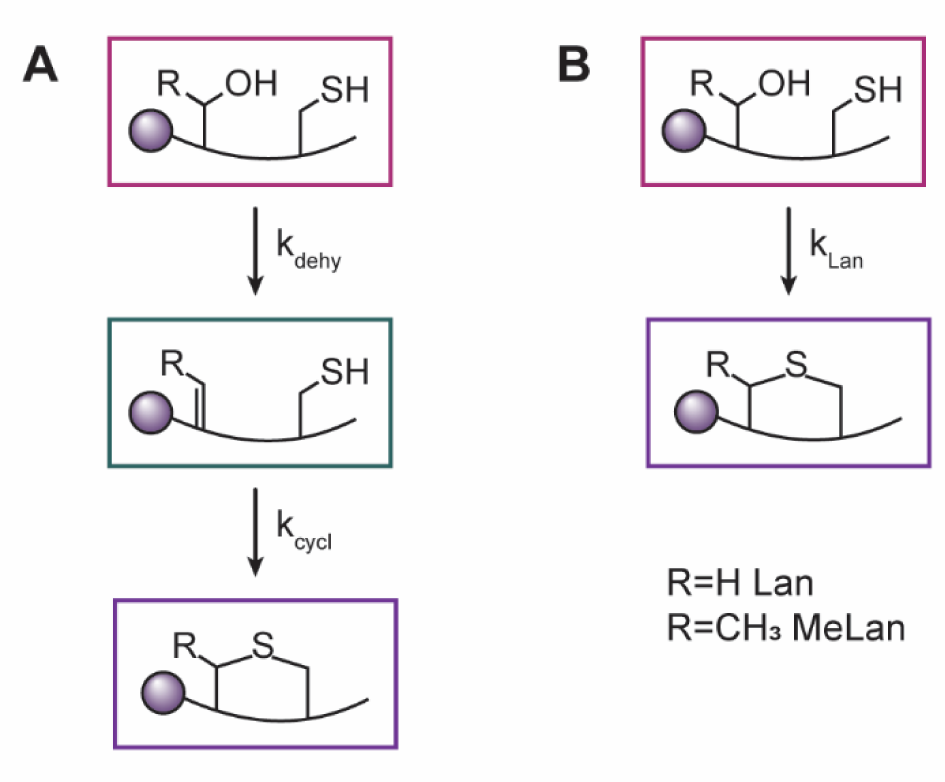
Proposed minimal kinetic models for isolated ring formation. Both models contain unmodified peptide (pink) and cyclized peptide (purple). **A)** Minimal kinetic model in which dehydrated peptide intermediate (teal) is present. This model was used for the formation of A, B, and B’ ring for WT ProcA3.3 and the A, A’ ring of the ProcA3.3 variant Z. **B)** Minimal kinetic model in which dehydrated intermediate was not present in sufficient quantity to use the former model with well-constrained values of rate constants. This model was used for the B’ ring of ProcA3.3 variant Z.

Analysis of individual rings in WT ProcA3.3 indicated that the rates for ring formation varied considerably (Table 2). The A ring, which is formed first in WT ProcA3.3, was formed fastest in the single ring variants with a cyclization rate of 20 min^-1^. The B’ ring, which is the C terminal non-overlapping ring, was the next fastest with a rate of 1.58 min^-1^, followed by the B ring which had the slowest measured rate of 0.53 min^-1^. These rate constants follow qualitative observations previously reported for WT ProcA3.3 that demonstrated that the A ring was formed first. Furthermore, when Cys14 involved in A ring formation was capped with a photolabile protecting group, the B’ ring was observed to form rather than the B ring.^29^ Thus, these individual ring cyclization rates are consistent with the experimentally observed ring cyclization order for WT ProcA3.3.

**Table 2.** Summary of Simulated Rate Constants for Isolated Lanthipeptide Rings.

| Isolated Ring | $k_x$ | Best fit ± SE (min<sup>-1</sup>) | FitSpace Boundaries (min<sup>-1</sup>)<sup>a</sup> | $k_{net}$ ($\mu\text{M}^{-1} \text{min}^{-1}$)<sup>**</sup> |
| --- | --- | --- | --- | --- |
| **ProcA3.3 WT** |  |  |  |  |
| A ring (Dhb) | $k_{dehy}$ | $9.1 \pm 0.8$ | 7.4 – 11.5 | 1.3 |
| | $k_{cycl}$ | $20 \pm 4$ | 14.1 – 37.1 | 2.1 |
| B ring (Dhb) | $k_{dehy}$ | $23 \pm 4$ | 15 – 50 | 2.3 |
| | $k_{cycl}$ | $0.53 \pm 0.03$ | 0.46 – 0.61 | 0.11 |
| B' ring (Dhb) | $k_{dehy}$ | $11 \pm 1$ | 8 – 15 | 1.5 |
| | $k_{cycl}$ | $1.58 \pm 0.09$ | 1.33 – 1.86 | 0.312 |
| **ProcA3.3 variant Z** |  |  |  |  |
| A ring (Dha) | $k_{dehy}$ | $8.3 \pm 0.5$ | 7.4 – 8.9 | 1.2 |
| | $k_{cycl}$ | $32 \pm 7$ | 28 – 60 | 2.8 |
| A' ring (Dha) | $k_{dehy}$ | $16 \pm 2$ | 14 – 18 | 1.9 |
| | $k_{cycl}$ | $120 \pm 90$ | 70 – 1090 | 3.7 |
| B' ring (Dha) | $k_{lan}$ | $89 \pm 31$ | 39 – 58 | 3.5 |
| A ring (Dhb) | $k_{dehy}$ | $6.9 \pm 0.5$ | 6.1 – 8.1 | 1.1 |
| | $k_{cycl}$ | $11 \pm 2$ | 8 – 18 | 1.5 |

<sup>a</sup>FitSpace boundaries are reported for a $\chi^2$ threshold boundary of 0.83 for ProcA3.3 WT and 0.96 for the ProcA3.3 variant. <sup>\*\*</sup> $k_{net} = (k_{on}k_x)/(k_{off} + k_x)$ . <sup>a</sup> $k_{on}$ is in units of $\mu\text{M}^{-1} \text{min}^{-1}$ . <sup>b</sup> $k_{on}$ and $k_{off}$ values were used from previously published data.<sup>30</sup>

Surprisingly, the isolated A’ ring was not efficiently cyclized by ProcM under these reaction conditions (Figure S10) even though this ring is formed in the presence of the B’ ring.^28^ This observation suggests that the presence of the B’ ring significantly accelerates the formation of the A’ ring within the WT ProcA3.3 sequence. Collectively, the cyclization rates of individual rings correctly predict that WT ProcA3.3 would generate an overlapping ring pattern.

### Single ring cyclization kinetics for ProcA3.3 variant Z

Single cyclization kinetics were also performed on peptides only able to form the A, A’ and B’ rings in the ProcA3.3 variant Z scaffold (Figures S11-13). The B ring variant was not tested in isolation for this scaffold because this ring has never been observed to form first in any fully modified ProcA3.3 peptide. Peaks corresponding to unmodified, dehydrated, and cyclized peptide species were observed and fractional abundance was calculated to monitor reaction progression. These data were used in the same minimal kinetic model as the ProcA3.3 WT isolated rings (Scheme 2A), except for the B’ ring of ProcA3.3 variant Z. For this substrate, the dehydrated intermediate was formed and consumed rapidly resulting in a fit to the model in Scheme 2A that did not yield well-constrained rate constants. In order to generate better constrained rate constants, a second model for individual cyclization rates was generated (Scheme 2B). In this model, the dehydrated intermediate was removed, and the dehydration and cyclization rates combined to give a lanthionine formation rate, *k_lan_*. In this model, *k_dehy_* and *k_cycl_* are coupled such that *k_lan_* would be representative of the rate limiting chemical modification, in this case the dehydration rate.

Similar to the observations with the ProcA3.3 WT scaffold, the rate of isolated lanthionine formation varied widely (Table 2). The rings present in the fully modified peptide, A’ and B’, were formed at rates that showed minimal buildup of the dehydrated intermediate, resulting in an inability to measure *k_cycl_* for the B’ ring as discussed above, and a large standard error for the A’ ring cyclization rate. Examining the rate boundaries using the FitSpace Explorer module of KinTek indicates that the A’ ring *k_cycl_* rate has a well constrained lower boundary (70 min^-1^) and a poorly defined upper boundary (1090 min^-1^). Therefore, it is not possible to directly compare the cyclization rates of the two rings. Given that the dehydration rate constant for the A’ ring (16 min^-1^) is less than that of the lanthionine formation rate for the B’ ring (89 min^-1^), and that the dehydration step is rate limiting for both ring formations, it is possible that the B’ ring in isolation is formed faster than the A’ ring in isolation.

Importantly, the A’ and B’ rings of the variant Z scaffold are formed at faster rates than the A ring in the variant Z scaffold (32 min^-1^). Therefore, the isolated ring rates correctly predict the final ring pattern for both ProcA3.3 WT and variant Z.

### Kinetics of lanthionine compared to methyllanthionine formation

Although many studies on lanthipeptides have shown that the biosynthetic machinery can accept Ser-to-Thr and Thr-to-Ser mutations leading to interchange of Lan and MeLan rings,^36–39^ these studies have typically been conducted on systems that have one lanthipeptide synthetase and one substrate. For these systems, substrate and enzymatic machinery have co-evolved under strong evolutionary pressure to make a bioactive product. Changing Lan to MeLan or vice-versa are minor changes (addition or removal of a methyl group) that are tolerated by these highly evolved biosynthetic systems. On the other hand, as has been discussed previously,^30,32^ the system under investigation here in which one enzyme has 30 different substrates cannot have evolved to very efficiently make a single product. Much evidence has been accumulated including the current study that the substrate sequence determines the outcome of catalysis.^32^ The experiments presented in this work show that interconversion of Lan to MeLan and vice-versa are not as straightforward for these systems and can lead to changes in the ring pattern formed. This finding raised a question that has not yet been addressed in lanthipeptide research: what is the difference in rate of formation of Lan and MeLan within the same sequence context? We felt that the single ring kinetics presented above provided an excellent opportunity to investigate this question. Thus, we performed kinetics for formation of the MeLan A ring in the variant Z scaffold (Figure S14) to provide the first direct examination of the rate changes in enzyme catalyzed cyclization of Lan versus MeLan. Similar to the other peptides, peaks corresponding to unmodified, dehydrated, and cyclized peptides were observed and fractional abundance was calculated to monitor reaction progression (Figure S14). The data were used in the same minimal kinetic model as the other isolated ring formations with a dehydrated intermediate (Scheme 2A). The data show that the rates of dehydration of Ser or Thr in the same sequence context were similar (8.3 vs 6.9 min^-1^; Table 2). However, cyclization of the ring containing Dhb (11 min^-1^, Table 2) was about three-fold slower than the Dha-containing counterpart (32 min^-1^), likely due to the increased steric hinderance and decreased electrophilicity.

## Conclusions

Kinetic analysis of ProcA3.3 WT and ProcA3.3 variant Z identified differences in both the order of modification and the rate of the post-translational modifications (PTMs) by ProcM that resulted in the change in ring pattern observed. Consistent with a prior kinetic study on ProcA2.8, the presence of PTMs altered the rate of modification within the peptide, with reactions slowing down as more modifications were present. Collectively, the current studies on the ProcA3.3 scaffolds indicate that both the identity of the dehydratable residues and residues not directly involved in ring formation can be important for the produced ring pattern. While the latter had been observed in previous studies,^23,32^ the notion that changing from Lan to MeLan (or vice versa) can change the ring pattern because of increased rates of formation of rings not present in the parent product is an important factor for predicting ring patterns, both for the products generated from novel peptide sequences in genomes and for library generation. Analysis of the rates of formation of isolated rings within the ProcA3.3 scaffolds indicate that the relative ring formation rates can be used to determine the final ring pattern, but cannot necessarily be used to determine the order of modifications in WT and variant Z scaffolds.

This study illustrates the complexity of ProcM catalyzed reactions, and that specific mechanistic knowledge is required to enable reliable methods for the prediction of final ring pattern on the basis of precursor peptide sequence. These findings raise questions whether RiPP biosynthetic enzymes that originate from biodiversity generating systems (i.e. with many physiological substrates having diverse sequences) are the most promising for generation of diverse engineered libraries. In the context of lanthipeptides, it may be that enzymes that have evolved to make single ring patterns robustly and more rapidly, but that still show high sequence tolerance in reliably generating that particular ring pattern may be better starting points.

## Acknowledgements

This work was supported by the National Institutes of Health (R37 GM058822 to W.A.V.). The Waters Q/ToF Synapt-G1 series mass spectrometer was purchased with a grant from the Howard Hughes Medical Institute (HHMI), and W.AV. is an HHMI Investigator.

