## Supporting Information for "Kinetic Analysis of Lanthipeptide Cyclization by Substrate-Tolerant ProcM"

### Supporting Information: Kinetic Analysis of Cyclization Reactions Performed by Substrate-Tolerant ProcM

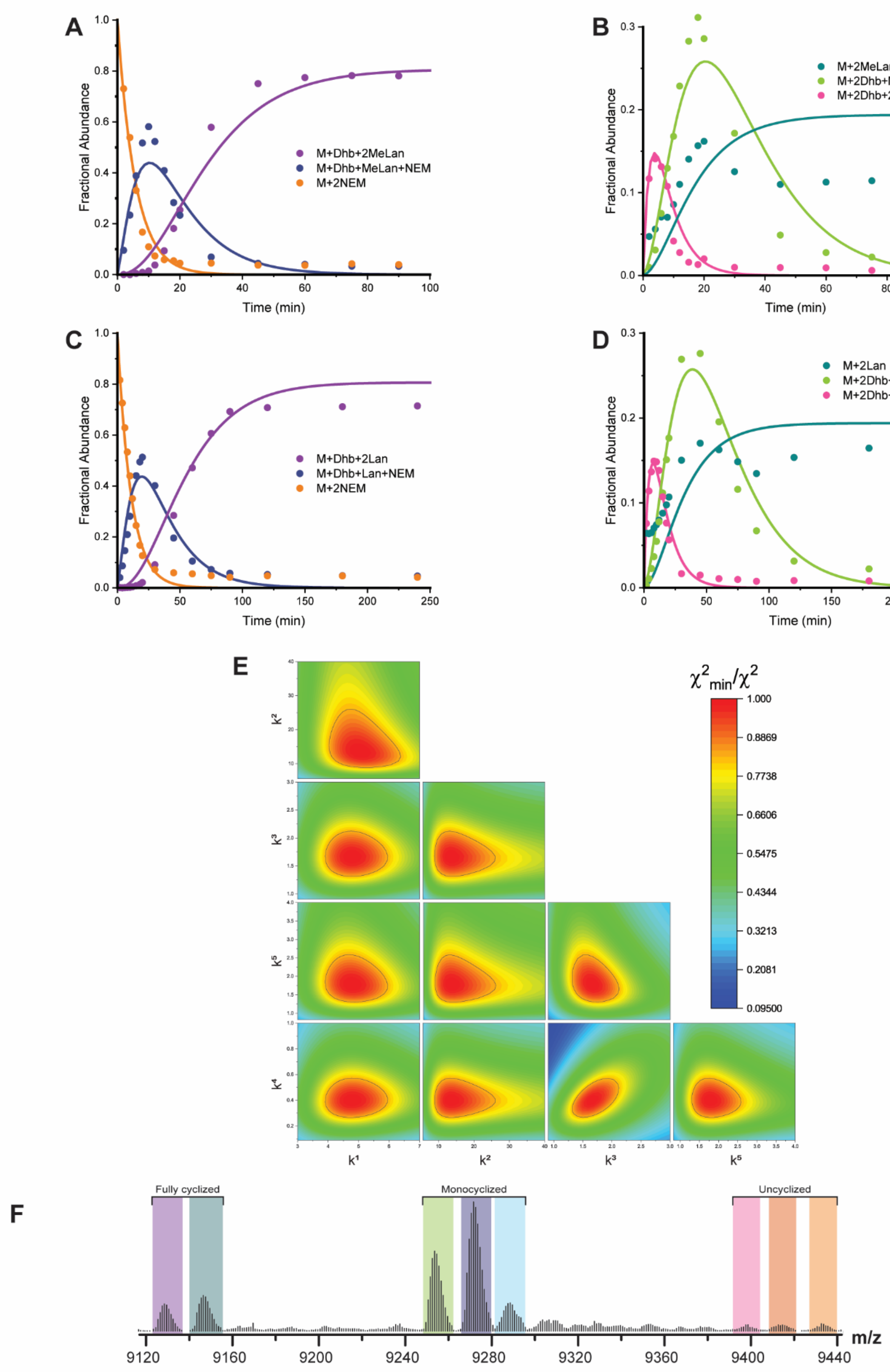
A
Fractional Abundance
Time (min)
M+Dhb+2MeLan
M+Dhb+MeLan+NEM
M+2NEM
B
Fractional Abundance
Time (min)
M+2MeLan
M+2Dhb+MeLan+NEM
M+2Dhb+2NEM
C
Fractional Abundance
Time (min)
M+Dhb+2Lan
M+Dhb+Lan+NEM
M+2NEM
D
Fractional Abundance
Time (min)
M+2Lan
M+2Dhb+Lan+NEM
M+2Dhb+2NEM
E
χ²min/χ²
1.000
0.8869
0.7738
0.6606
0.5475
0.4344
0.3213
0.2081
0.09500
F
Fully cyclized
Monocyclized
Uncyclized
m/z
9120
9160
9200
9240
9280
9320
9360
9400
9440

**Figure S1. ProcA3.3 WT kinetics.** Species observed include unmodified peptide (orange), peptide containing one MeLan ring and one dehydration (dark blue), the final product (purple), uncyclized peptide with two dehydrations (pink), peptide containing one MeLan ring with two dehydrations (green), and peptide containing two MeLan rings with no dehydrations (cyan). **A/B)** Time course for species in a reaction containing 40 μM peptide and 2 μM ProcM. **C/D)** Time course for species for reaction containing 80 μM peptide and 2 μM ProcM. **E)** Confidence contours used to evaluate the upper and lower bounds for each rate constant listed in Table 1 of the main text. The kinetic data shown here and in Figure 3 of the main text were simulated using the kinetic model shown in Scheme 1A. The fit of the rate constants listed in Table 1 yielded a $\chi_{min}^2$ of 131.8 ($\chi^2$/DOF = 0.438). The confidence contours indicate the $\chi_{min}^2/\ \chi^2$ value as a function of variable parameters indicated on the x and y axis. Values with a $\chi_{min}^2/\ \chi^2$ value greater than 0.833 are within the black line indicated on the plot. This is the cut off value that was utilized for all ProcA3.3 WT reactions. **F)** Representative deconvoluted ESI mass spectra of ProcA3.3 WT reaction with ProcM at 20 minutes of the 60 μM reaction. Data has been represented in centroid mode to show monoisotopic species as opposed to profile mode as shown in Figure 3A. See Table S1 for a full listing of calcuylated and observed m/z values for all ions. For each trace, the y-axis is scaled to the intensity of the highest peak present. The sample was treated with NEM to interrogate the cyclization state of the Cys residues. The coloring of each ion matches those in other panels and those in Figure 3.

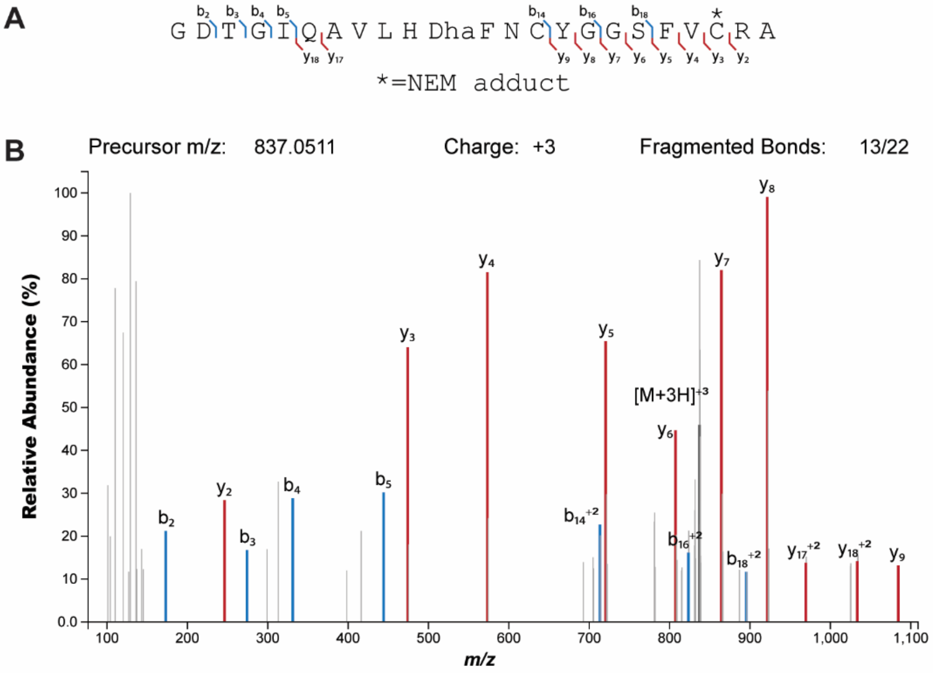

C

| Ion | Calc. mass | Obs. Mass | Error (ppm) |
|---|---|---|---|
| $b_2^+$ | 173.0557 | 173.0557 | 0.1 |
| $y_2^+$ | 246.1563 | 246.1561 | 1.0 |
| $b_3^+$ | 274.1054 | 274.1034 | 7.4 |
| $b_4^+$ | 331.1248 | 331.1248 | -0.1 |
| $b_5^+$ | 444.2091 | 444.2089 | 0.5 |
| $y_3^+$ | 474.2121 | 474.213 | -1.8 |
| $y_4^+$ | 573.2797 | 573.2814 | -2.9 |
| $b_{14}^{+2}$ | 713.3316 | 713.3326 | -1.4 |
| $y_5^+$ | 720.3483 | 720.3498 | -2.1 |
| $y_6^+$ | 807.3811 | 807.3818 | -0.9 |
| $b_{16}^{+2}$ | 823.3737 | 823.375 | -1.6 |
| $[M+3H]^{+3}$ | 837.0498 | 837.0511 | -1.6 |
| $y_7^+$ | 864.4013 | 864.4033 | -2.3 |
| $b_{18}^{+2}$ | 895.3987 | 895.4018 | -3.4 |
| $y_8^+$ | 921.4233 | 921.4247 | -1.6 |
| $y_{17}^{+2}$ | 969.4404 | 969.4429 | -2.6 |
| $y_{18}^{+2}$ | 1033.4737 | 1033.4722 | 1.4 |
| $y_9^+$ | 1084.4844 | 1084.4881 | -3.4 |

**Figure S2. Fragmentation analysis of singly cyclized intermediate for ProcA3.3 variant Z**. **A)** Sequence of ProcA3.3 variant singly cyclized intermediate after peptide cleavage. Observed fragmentation is indicated with labeled y and b ions. **B)** Tandem-MS fragmentation of spectrum with corresponding peaks labeled. **C)** Plot of observed errors for each peak calculated in ppm. Panels A and B are modified from those produced on Interactive Peptide Spectral Annotator.[1]

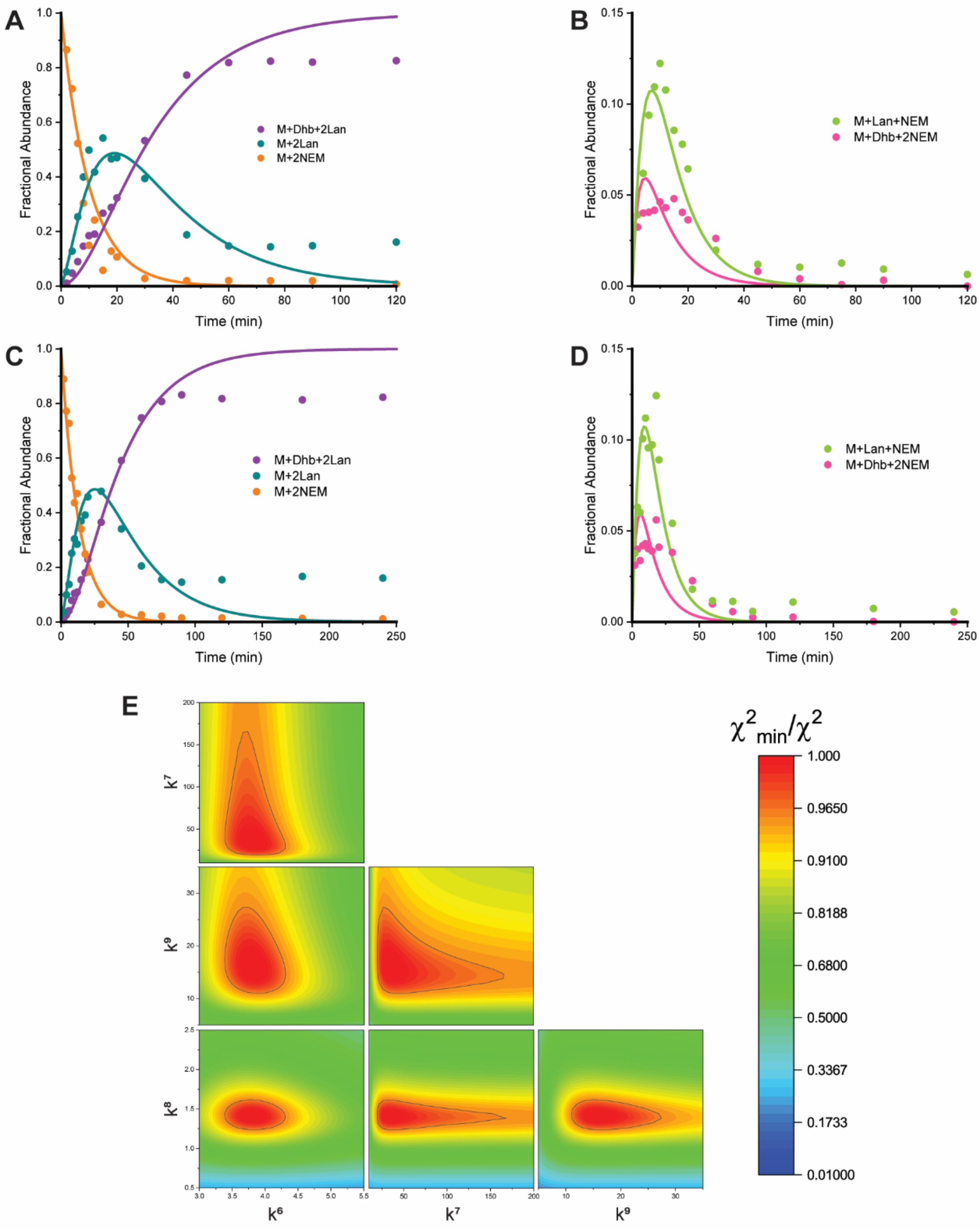

**Figure S3. ProcA3.3 Variant Z kinetics.** Species observed include unmodified peptide (orange), peptide containing two Lan rings with no additional dehydrations (cyan), the final product (purple), uncyclized peptide with one dehydration (pink), and peptide containing one Lan ring (green). **A/B)** Time course for species for the reaction containing 60 µM peptide and 2 µM ProcM. **C/D)** Time course for the reaction containing 80 µM peptide and 2 µM ProcM. **E)** Confidence contours used to evaluate the upper and lower bounds for each rate constant listed in Table 1 of the main text.

The kinetic data shown here and in Figure 4 of the main text were simulated using the kinetic model shown in Scheme 1B. The fit of the rate constants listed in Table 1 yielded a $\chi_{min}^2$ of 394.7 ($\chi^2$/DOF = 1.573). The confidence contours indicate the $\chi_{min}^2/\ \chi^2$ value as a function of variable parameters indicated on the x and y axis. Values with a $\chi_{min}^2/\ \chi^2$ value greater than 0.96 are within the black line indicated on the plot. This is the cut off value that was utilized for all ProcA3.3 variants. All rate constants within Scheme 1B were well-defined by the data collected, however the $k_7$ rate constants have well defined lower boundaries, with less well-defined upper boundaries.

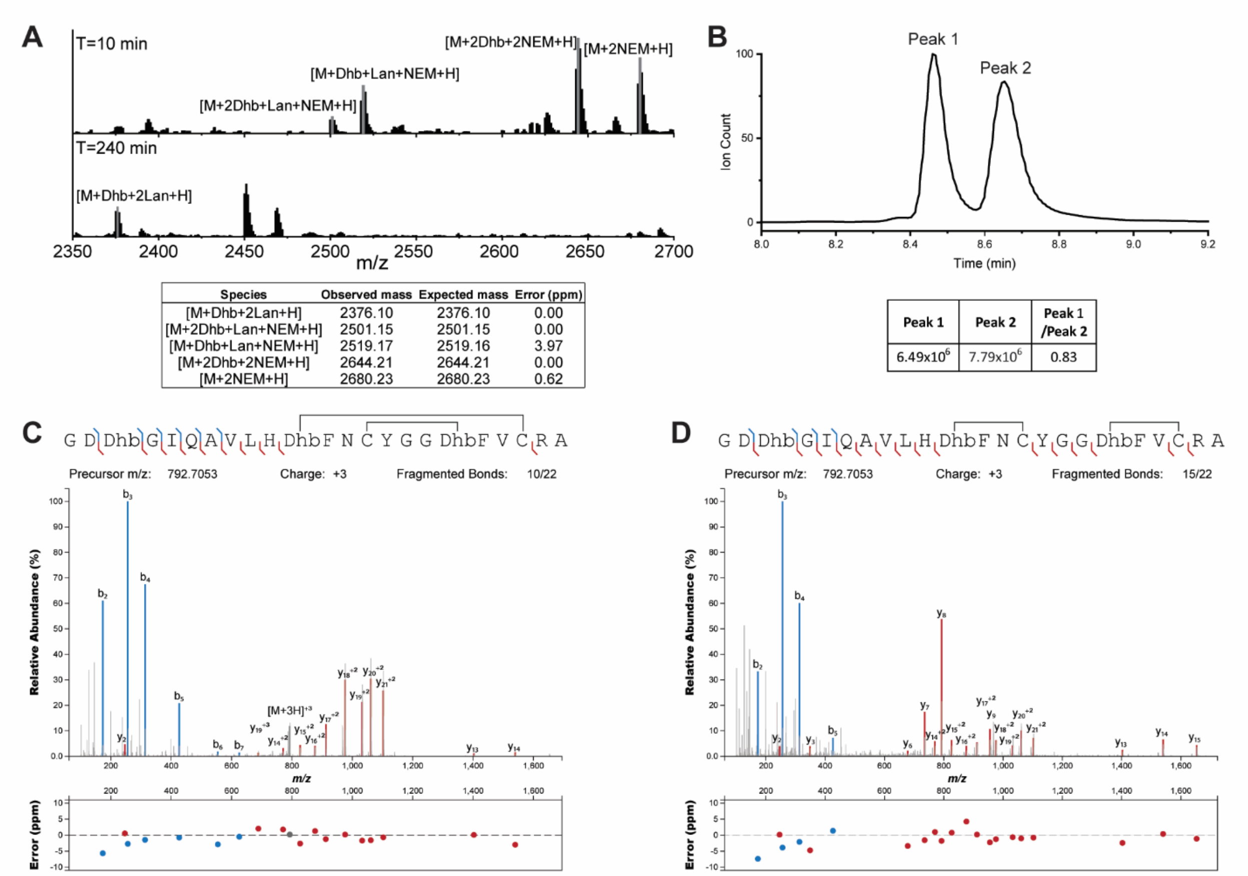

**Figure S4. MeLan-forming ProcA3.3 Variant Z analysis. A)** Deconvoluted ESI mass spectra of MeLan-forming ProcA3.3 variant Z modified by ProcM at the 10- and 240-minute time points. The sample was treated with NEM to interrogate the cyclization state of the Cys residues. Peaks corresponding to peptide intermediates are labeled, and observed and expected masses are provided below the spectrum. **B)** Extracted ion chromatogram of MeLan containing ProcA3.3 variant Z products (m/z=792.7054, z=3) after an overnight reaction. **C/D)** Observed fragmentation is indicated with labeled y and b ions of both peptides associated with peaks 1 and 2 of the extracted ion chromatogram, respectively. Error is depicted below for each corresponding labeled peak above. Panels C and D were produced using Interactive Peptide Spectral Annotator.[1] The fragmentation observed indicates that peak 1 corresponds to product with an overlapping ring pattern and peak 2 corresponds to product with a non-overlapping ring pattern. Samples for all panels were treated with LahT$_{150}$,[2] and protein was precipitated via MeOH addition prior to analysis (see methods section).

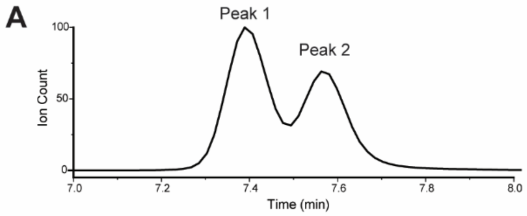

| Peak 1 | Peak 2 | Peak 1 /Peak 2 |
|---|---|---|
| 8.09x10$^5$ | 6.02x10$^5$ | 1.34 |

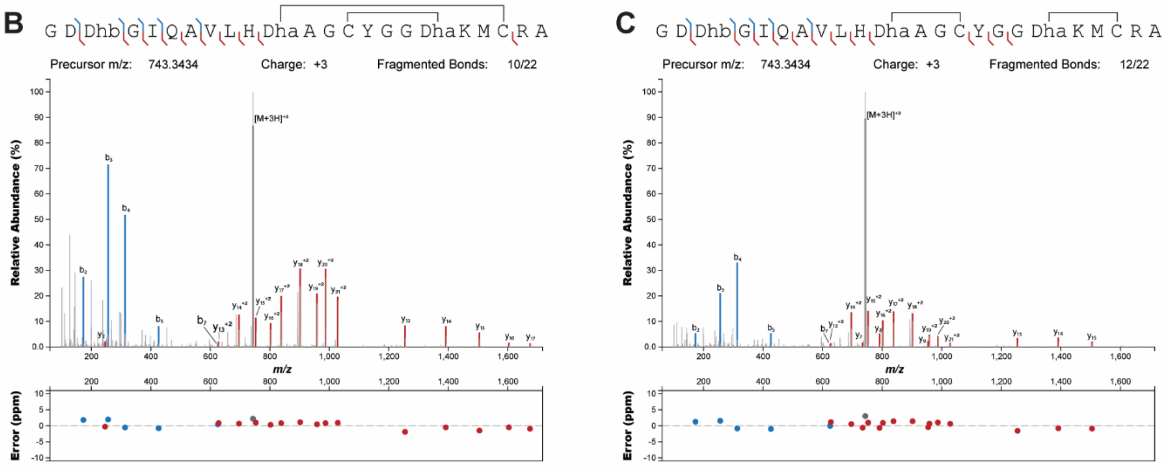

**Figure S5. Lan-forming ProcA3.3 WT analysis. A)** Extracted ion chromatogram of products formed from Lan-forming ProcA3.3 WT (m/z=743.3435, z=3) after the conclusion of a four-hour reaction. **B/C)** Observed fragmentation is indicated with labeled y and b ions of both Peak 1 and 2 of the extracted ion chromatogram, respectively. Error is depicted below for each corresponding labeled peak above. Panels B and C were produced using Interactive Peptide Spectral Annotator.[1] Fragmentation indicates that peak 1 corresponds to the overlapping ring pattern and peak 2 corresponds to the non-overlapping ring pattern. Samples for all panels were treated with $LahT_{150}$, and protein was precipitated via MeOH addition prior to analysis (see methods section).

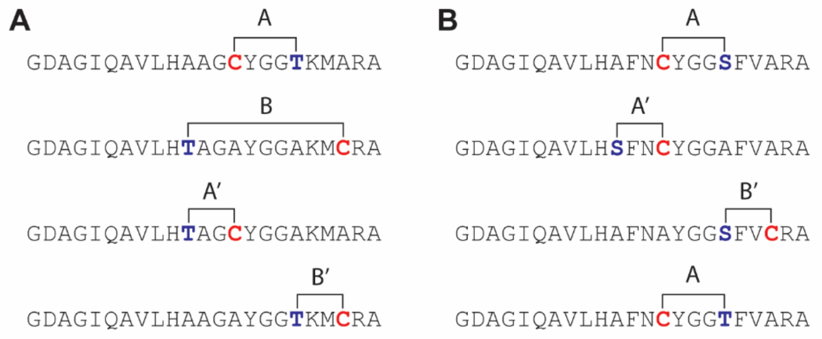

**Figure S6. ProcA3.3 peptides used for isolated ring kinetics**. **A)** The four variants used for kinetic analysis of individual cyclization rates on the ProcA3.3 WT scaffold. **B)** The four variants used for kinetic analysis of individual cyclization rates on the ProcA3.3 variant Z scaffold. A peptide that would form the B ring was not utilized for kinetic studies for this variant scaffold since this ring is never formed first.

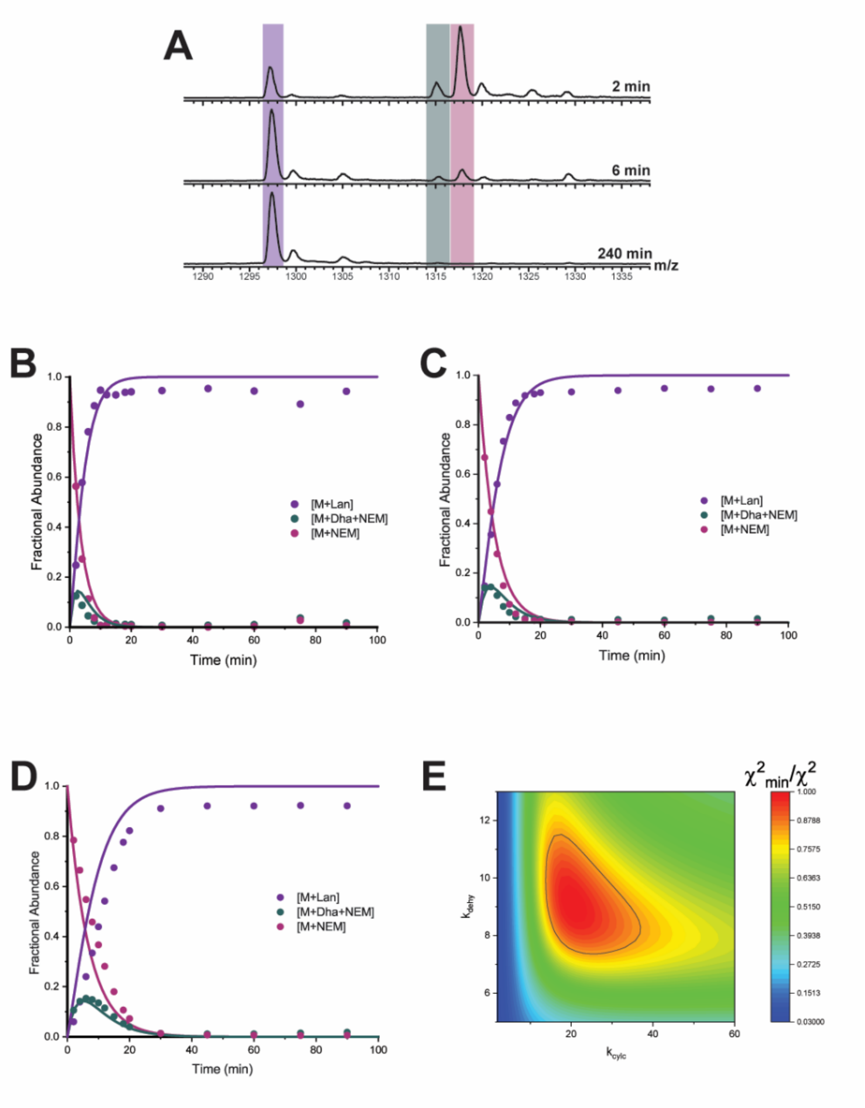

**Figure S7. ProcA3.3 A ring kinetics. A)** Representative ESI mass spectra of the 8+ charge state at multiple time points. For each trace, the y-axis is scaled to the intensity of the highest peak present. The sample was treated with NEM to interrogate the cyclization state of the Cys residue. Species observed include unmodified (pink), dehydrated (teal), and cyclized (purple) peptide. **B)** Time course for reaction containing 40 µM peptide and 2 µM ProcM. **C)** Time course for reaction containing 60 µM peptide and 2 µM ProcM. **D)** Time course for reaction containing 80 µM peptide and 2 µM ProcM. **E)** Confidence contour for the cyclization of the ProcA3.3 A ring in isolation was calculated with rate constants listed in Table 2 of main text. The kinetic data shown here were simulated using the kinetic model shown in Scheme 2A of the main text. The fit of the rate constants listed in Table 2 yielded a $\chi_{min}^2$ of 1813.9 ($\chi^2$/DOF = 12.3). The confidence contours indicate the $\chi_{min}^2/\chi^2$ value as a function of variable parameters indicated on the x and y axis. Values with a $\chi_{min}^2/\chi^2$ value greater than 0.833 are within the black line indicated on the plot.

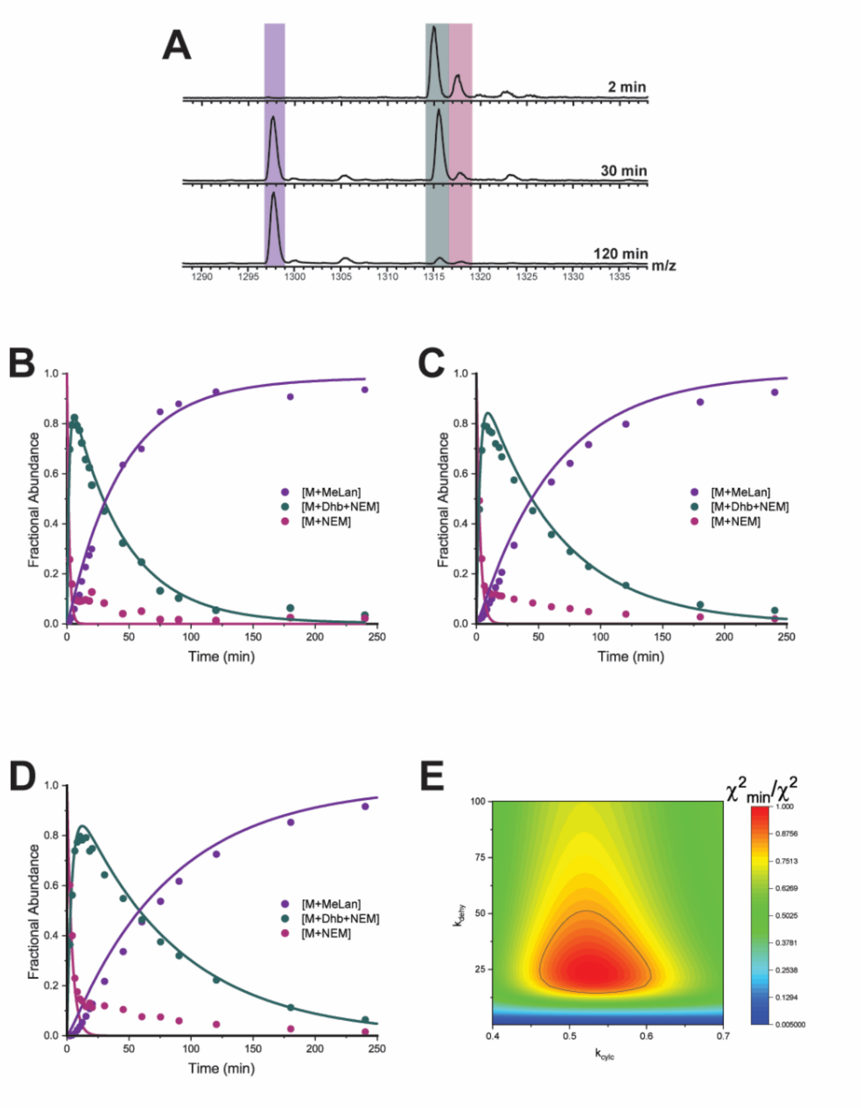

**Figure S8. ProcA3.3 B ring kinetics. A)** Representative ESI mass spectra of the 8+ charge state at multiple time points. For each trace, the y-axis is scaled to the intensity of the highest peak present. The sample was treated with NEM to interrogate the cyclization state of the Cys residue. Species observed include unmodified (pink), dehydrated (teal), and cyclized (purple) peptide. **B)** Time course for reaction containing 40 μM peptide and 2 μM ProcM. **C)** Time course for reaction containing 60 μM peptide and 2 μM ProcM. **D)** Time course for reaction containing 80 μM peptide and 2 μM ProcM. **E)** Confidence contour for the cyclization of the ProcA3.3 B ring in isolation was calculated with rate constants listed in Table 2 of main text. The kinetic data shown here were simulated using the kinetic model shown in Scheme 2A of the main text. The fit of the rate constants listed in Table 2 yielded a $\chi_{min}^2$ of 800.9 ($\chi^2$/DOF = 5.3). The confidence contours indicate the $\chi_{min}^2/\chi^2$ value as a function of variable parameters indicated on the x and y axis. Values with a $\chi_{min}^2/\chi^2$ value greater than 0.833 are within the black line indicated on the plot.

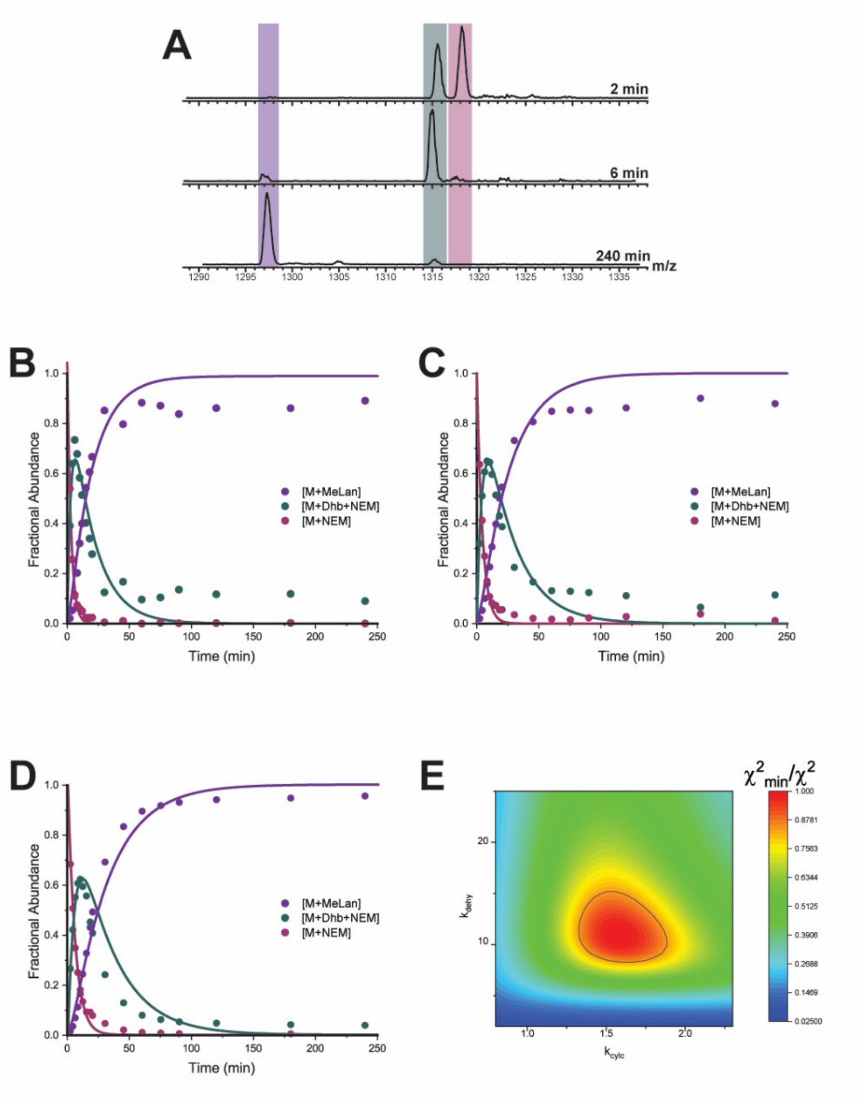

**Figure S9. ProcA3.3 B' ring kinetics. A)** Representative ESI mass spectra of the 8+ charge state at multiple time points. For each trace, the y-axis is scaled to the intensity of the highest peak present. The sample was treated with NEM to interrogate the cyclization state of the Cys residue. Species observed include unmodified (pink), dehydrated (teal), and cyclized (purple) peptide. **B)** Time course for reaction containing 40 µM peptide and 2 µM ProcM. **C)** Time course for reaction containing 60 µM peptide and 2 µM ProcM. **D)** Time course for reaction containing 80 µM peptide and 2 µM ProcM. **E)** Confidence contour for the cyclization of the ProcA3.3 B' ring in isolation was calculated with rate constants listed in Table 2 of main text. The fit of the rate constants listed in Table 2 yielded a $\chi_{min}^2$ of 396 ($\chi^2$/DOF = 2.6). The kinetic data shown here were simulated using the kinetic model shown in Scheme 2A of the main text. The confidence contours indicate the $\chi_{min}^2/\chi^2$ value as a function of variable parameters indicated on the x and y axis. Values with a $\chi_{min}^2/\chi^2$ value greater than 0.833 are within the black line indicated on the plot.

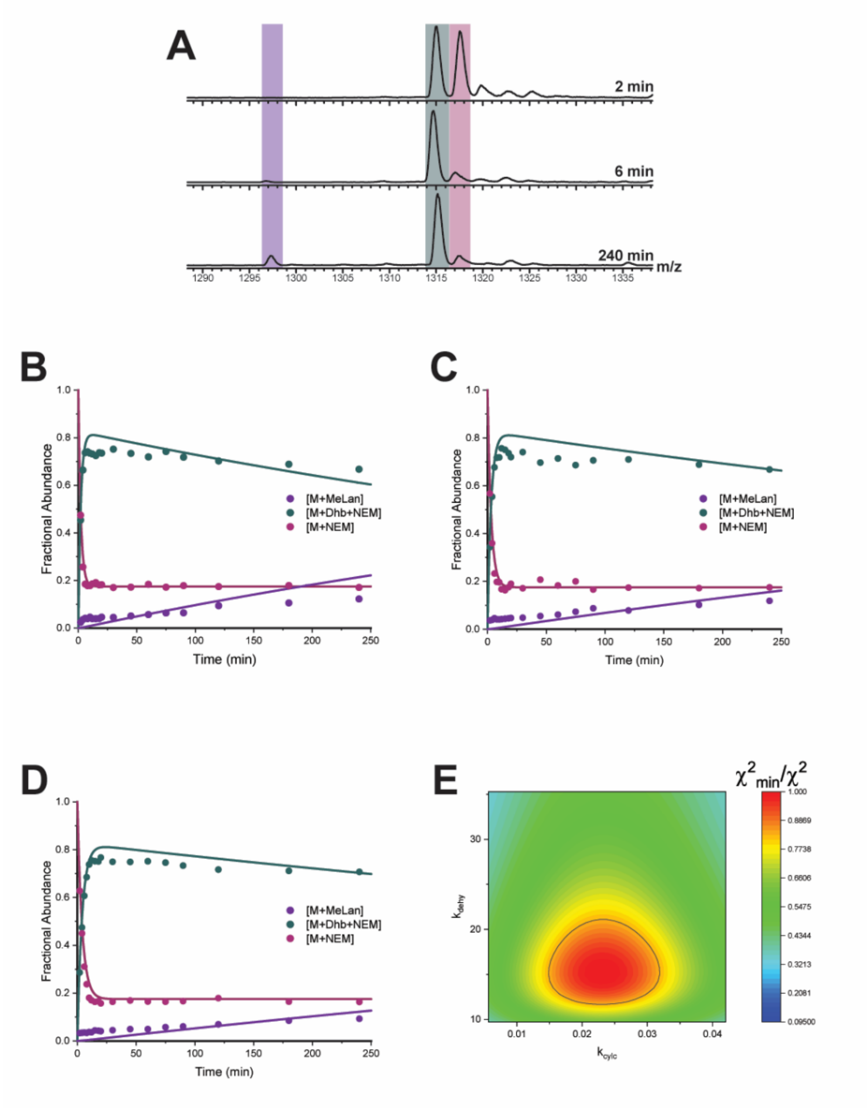

**Figure S10. ProcA3.3 A' ring kinetics. A)** Representative ESI mass spectra of the 8+ charge state at multiple time points. For each trace, the y-axis is scaled to the intensity of the highest peak present. The sample was treated with NEM to interrogate the cyclization state of the Cys residue. Species observed include unmodified (pink), dehydrated (teal), and cyclized (purple) peptide. This data was unable to be modeled due to the lack of kinetically competent cyclization. **B)** Time course for reaction containing 40 µM peptide and 2 µM ProcM. **C)** Time course for reaction containing 60 µM peptide and 2 µM ProcM. **D)** Time course for reaction containing 80 µM peptide and 2 µM ProcM. **E)** Confidence contour for the cyclization of the ProcA3.3 A' ring in isolation. The fit of the rate constants yielded a $\chi_{min}^2$ of 1231 ($\chi^2$/DOF = 8.2). The kinetic data shown here were simulated using the kinetic model shown in Scheme 2A of the main text. The confidence contours indicate the $\chi_{min}^2/\chi^2$ value as a function of variable parameters indicated on the x and y axis. Values with a $\chi_{min}^2/\chi^2$ value greater than 0.833 are within the black line indicated on the plot.

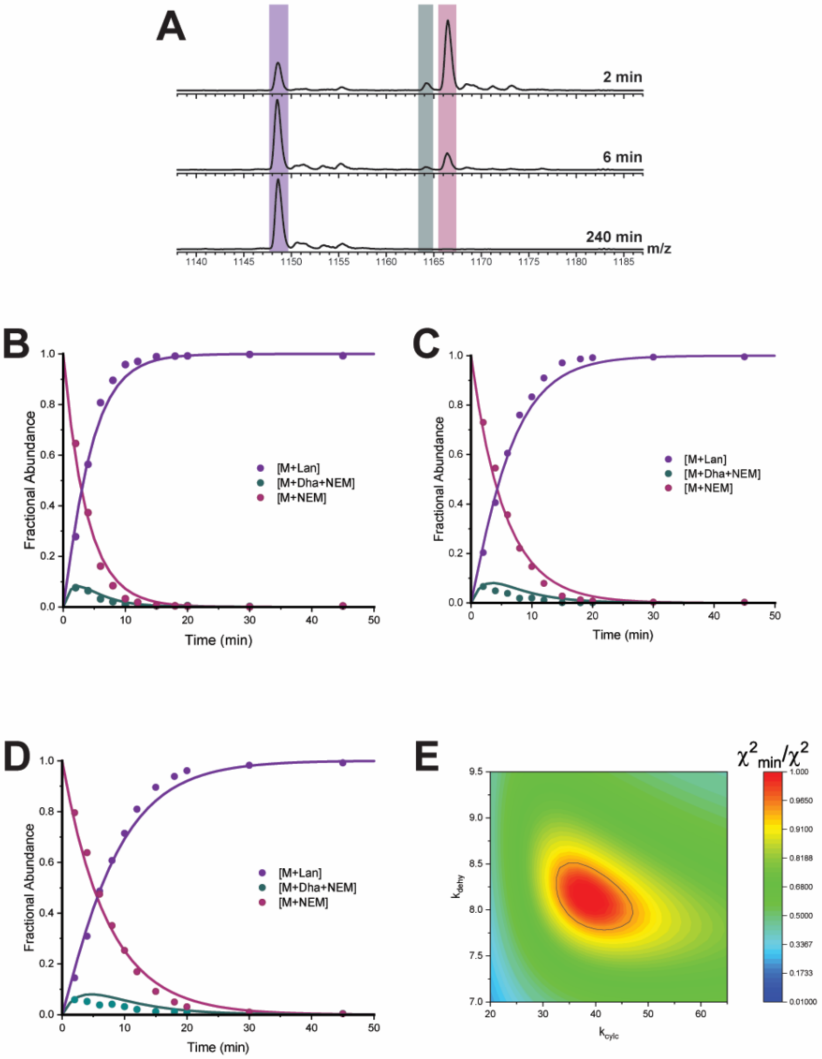

**Figure S11. ProcA3.3 Variant Z A ring kinetics. A)** Representative ESI mass spectra of the 8+ charge state at multiple time points. For each trace, the y-axis is scaled to the intensity of the highest peak present. The sample was treated with NEM to interrogate the cyclization state of the Cys residue. Species observed include unmodified (pink), dehydrated (teal), and cyclized (purple) peptide. **B)** Time course for reaction containing 40 µM peptide and 2 µM ProcM. **C)** Time course for reaction containing 60 µM peptide and 2 µM ProcM. **D)** Time course for reaction containing 80 µM peptide and 2 µM ProcM. **E)** Confidence contour for the cyclization of the ProcA3.3 variant Z A ring in isolation was calculated with rate constants listed in Table 2 of main text. The kinetic data shown here were simulated using the kinetic model shown in Scheme 2A of the main text. The fit of the rate constants listed in Table 2 yielded a $\chi_{min}^2$ of 787.2 ($\chi^2$/DOF = 5.7). The confidence contours indicate the $\chi_{min}^2/\ \chi^2$ value as a function of variable parameters indicated on the x and y axis. Values with a $\chi_{min}^2/\ \chi^2$ value greater than 0.96 are within the black line indicated on the plot.

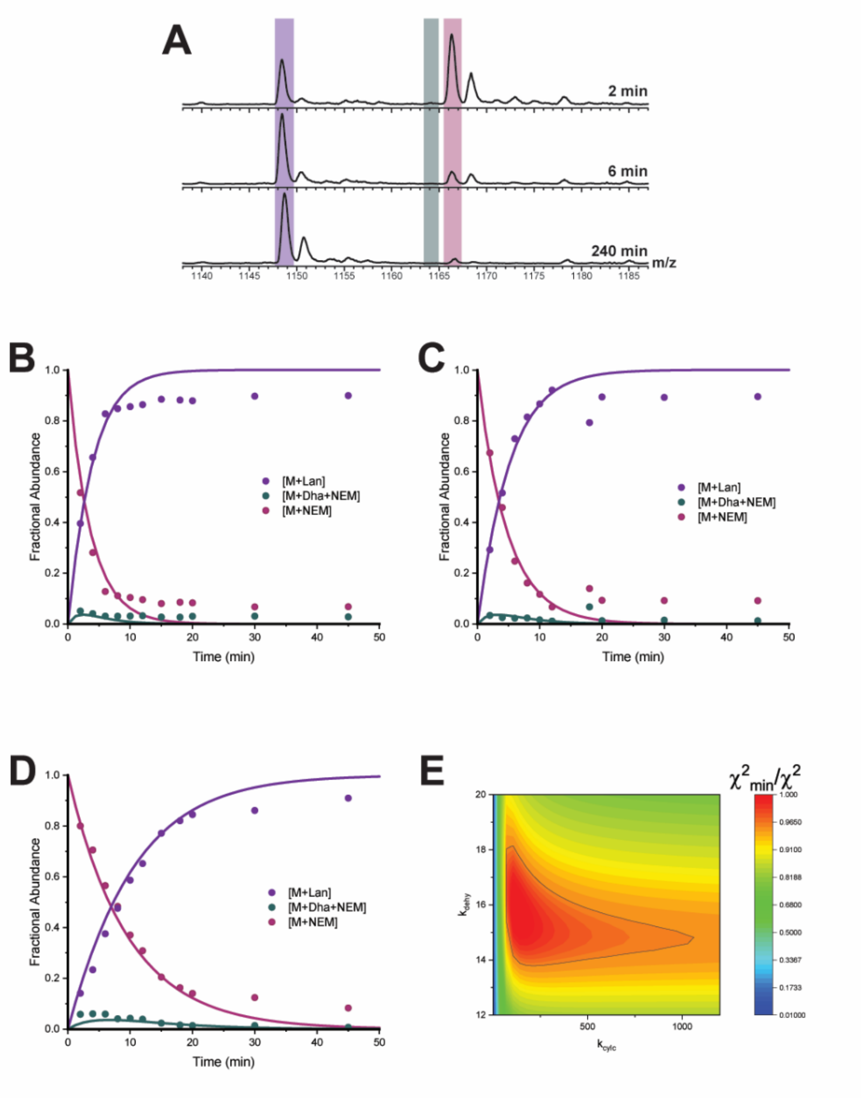

**Figure S12. ProcA3.3 Variant Z A' ring kinetics. A)** Representative ESI mass spectra of the 8+ charge state at multiple time points. For each trace, the y-axis is scaled to the intensity of the highest peak present. The sample was treated with NEM to interrogate the cyclization state of the Cys residue. Species observed include unmodified (pink), dehydrated (teal), and cyclized (purple) peptide. **B)** Time course for reaction containing 60 µM peptide and 2 µM ProcM. **C)** Time course for reaction containing 80 µM peptide and 2 µM ProcM. **D)** Time course for reaction containing 80 µM peptide and 1 µM ProcM. **E)** Confidence contour for the cyclization of the ProcA3.3 variant Z A' ring in isolation was calculated with rate constants listed in Table 2 of main text. The kinetic data shown here were simulated using the kinetic model shown in Scheme 2A of the main text. The fit of the rate constants listed in Table 2 yielded a $\chi_{min}^2$ of 1779.6 ($\chi^2$/DOF = 12.3). The confidence contours indicate the $\chi_{min}^2$/ $\chi^2$ value as a function of variable parameters indicated on the x and y axis. Values with a $\chi_{min}^2$/ $\chi^2$ value greater than 0.96 are within the black line indicated on the plot.

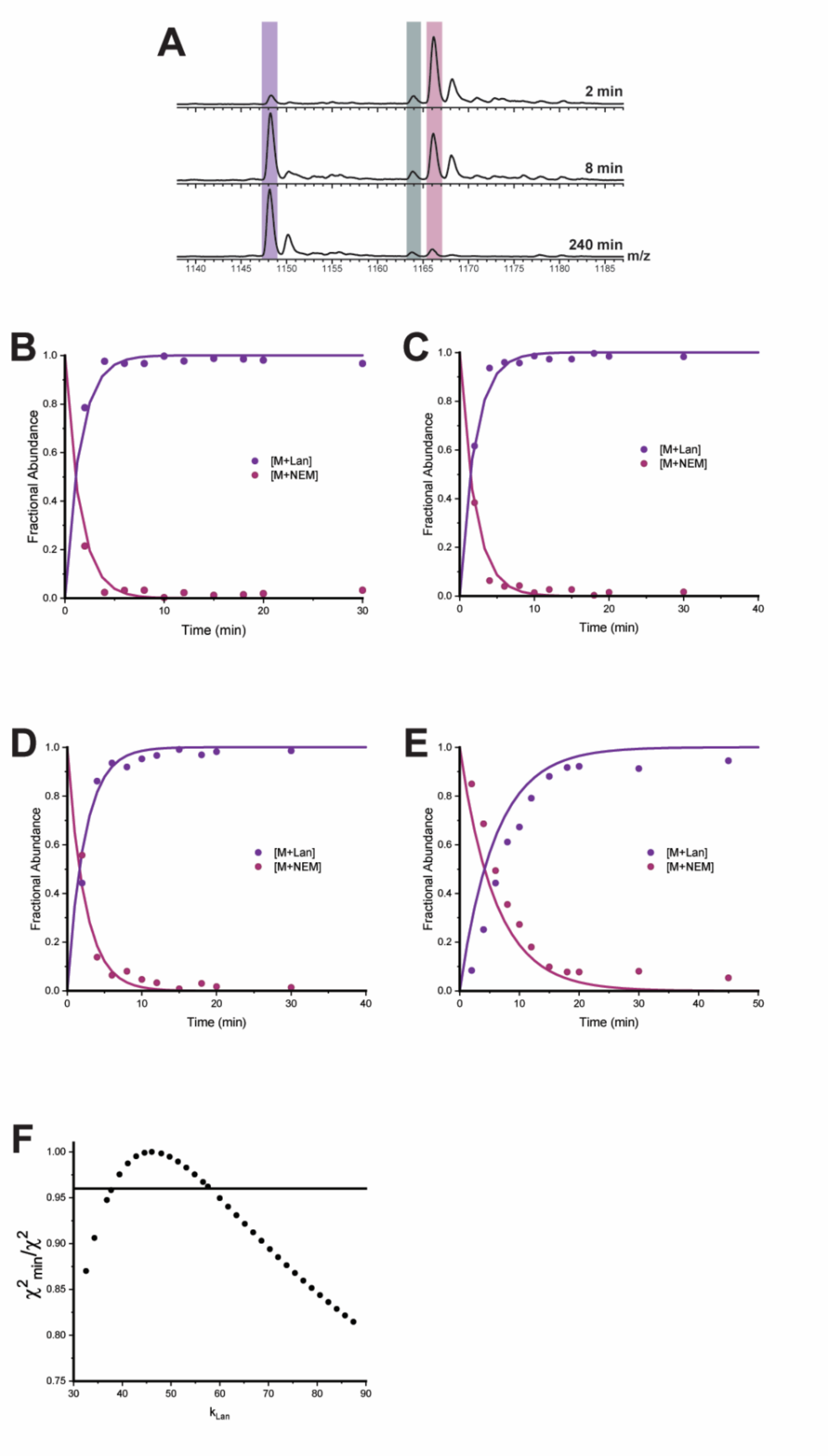

**Figure S13. ProcA3.3 Variant Z B' ring kinetics. A)** Representative ESI mass spectra of the 8+ charge state at multiple time points. For each trace, the y-axis is scaled to the intensity of the highest peak present. The sample was treated with NEM to interrogate the cyclization state of the Cys residue. Species observed include unmodified (pink) and cyclized (purple) peptide. **B)** Time course for reaction containing 40 μM peptide and 2 μM ProcM. **C)** Time course for reaction containing 60 μM peptide and 2 μM ProcM. **D)** Time course for reaction containing 80 μM peptide and 2 μM ProcM. **E)** Time course for reaction containing 80 μM peptide and 1 μM ProcM. **F)** One-dimensional confidence contour for the cyclization of the ProcA3.3 variant Z B' ring in isolation was calculated with the $k_{lan}$ rate constant listed in Table 2 of the main text. The kinetic data shown here were simulated using the kinetic model shown in Scheme 2B of the main text. The fit of the rate constants listed in Table 2 yielded a $\chi_{min}^2$ of 1944.5 ($\chi^2$/DOF = 5.4). The confidence contours indicate the $\chi_{min}^2/\ \chi^2$ value as a function of $k_{lan}$. Values with a $\chi_{min}^2/\ \chi^2$ value greater than 0.96 are above the black line indicated on the plot.

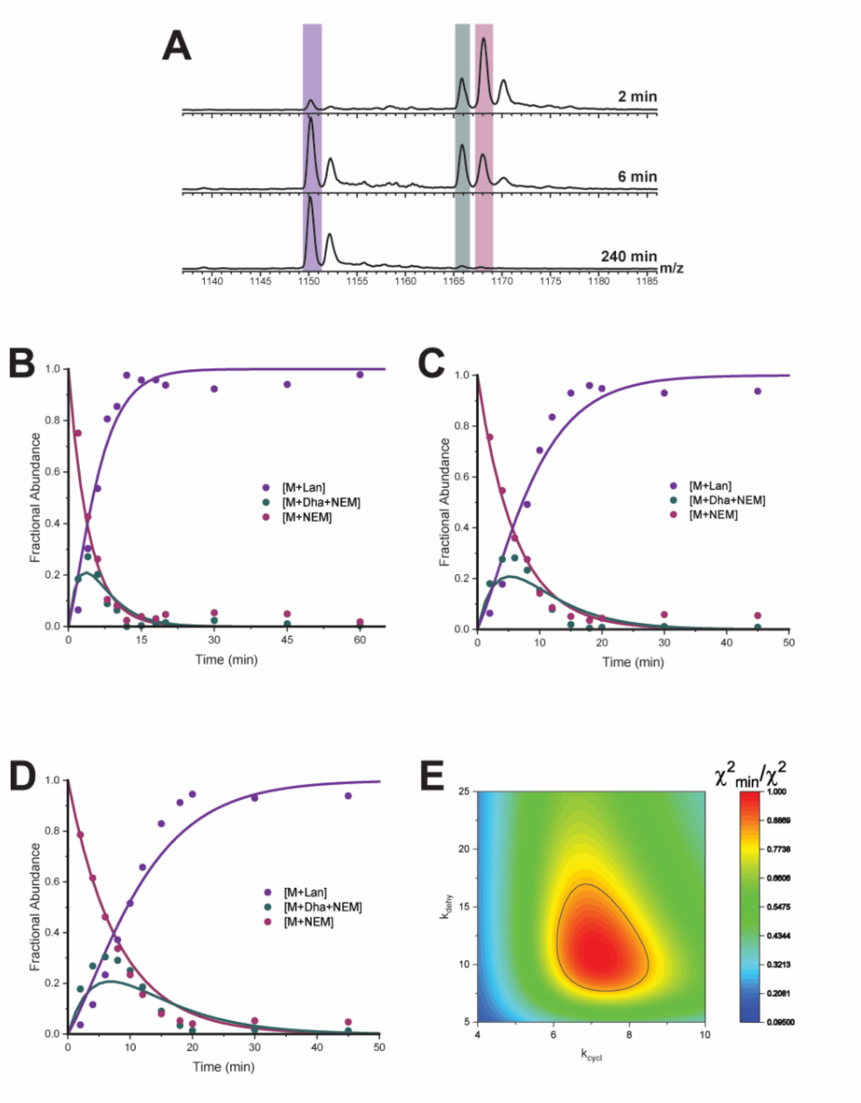

**Figure S14. ProcA3.3 Variant Z MeLan A ring kinetics. A)** Representative ESI mass spectra of the 8+ charge state at multiple time points. For each trace, the y-axis is scaled to the intensity of the highest peak present. The sample was treated with NEM to interrogate the cyclization state of the Cys residue. Species observed include unmodified (pink), dehydrated (teal), and cyclized (purple) peptide. **B)** Time course for reaction containing 40 µM peptide and 2 µM ProcM. **C)** Time course for reaction containing 60 µM peptide and 2 µM ProcM. **D)** Time course for reaction containing 80 µM peptide and 2 µM ProcM. **E)** Confidence contour for the cyclization of the ProcA3.3 variant Z MeLan A ring in isolation was calculated with rate constants listed in Table 2 of main text. The kinetic data shown here were simulated using the kinetic model shown in Scheme 2A of the main text. The fit of the rate constants listed in Table 2 yielded a $\chi_{min}^2$ of 325.9 ($\chi^2$/DOF = 2.2). The confidence contours indicate the $\chi_{min}^2$/ $\chi^2$ value as a function of variable parameters indicated on the x and y axis. Values with a $\chi_{min}^2$/ $\chi^2$ value greater than 0.83 are within the black line indicated on the plot.

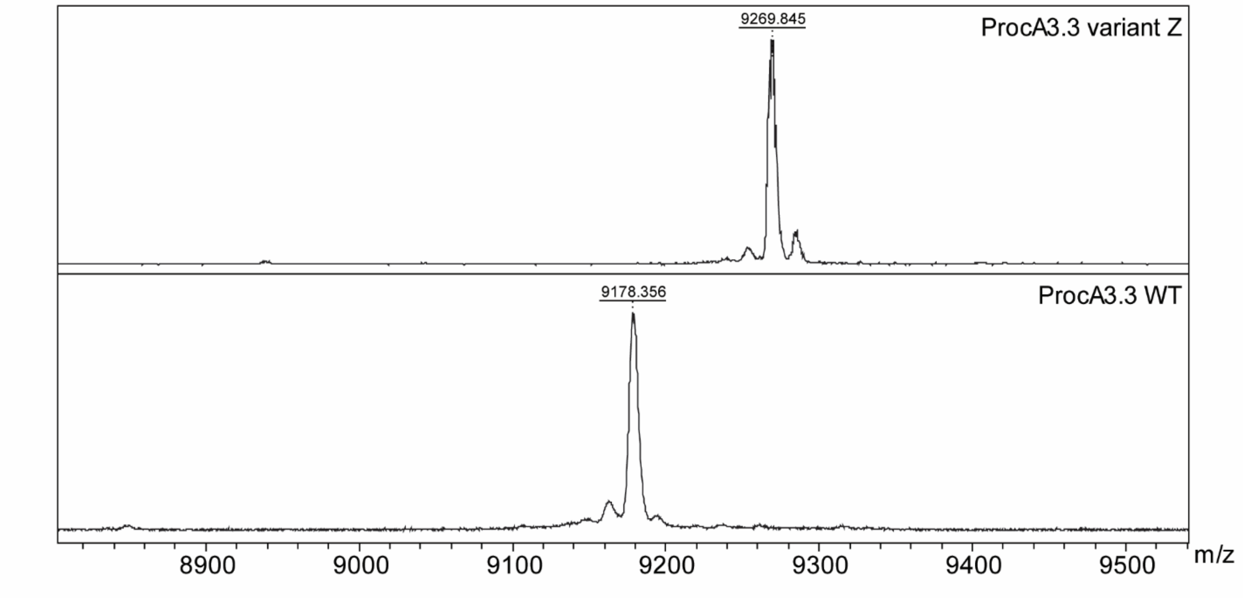

**Figure S15. MALDI-ToF mass spectra for parent ProcA3.3 peptides**. MALDI-ToF mass spectra were taken after peptide purification, thrombin digestion, and HPLC purification of the peptides.

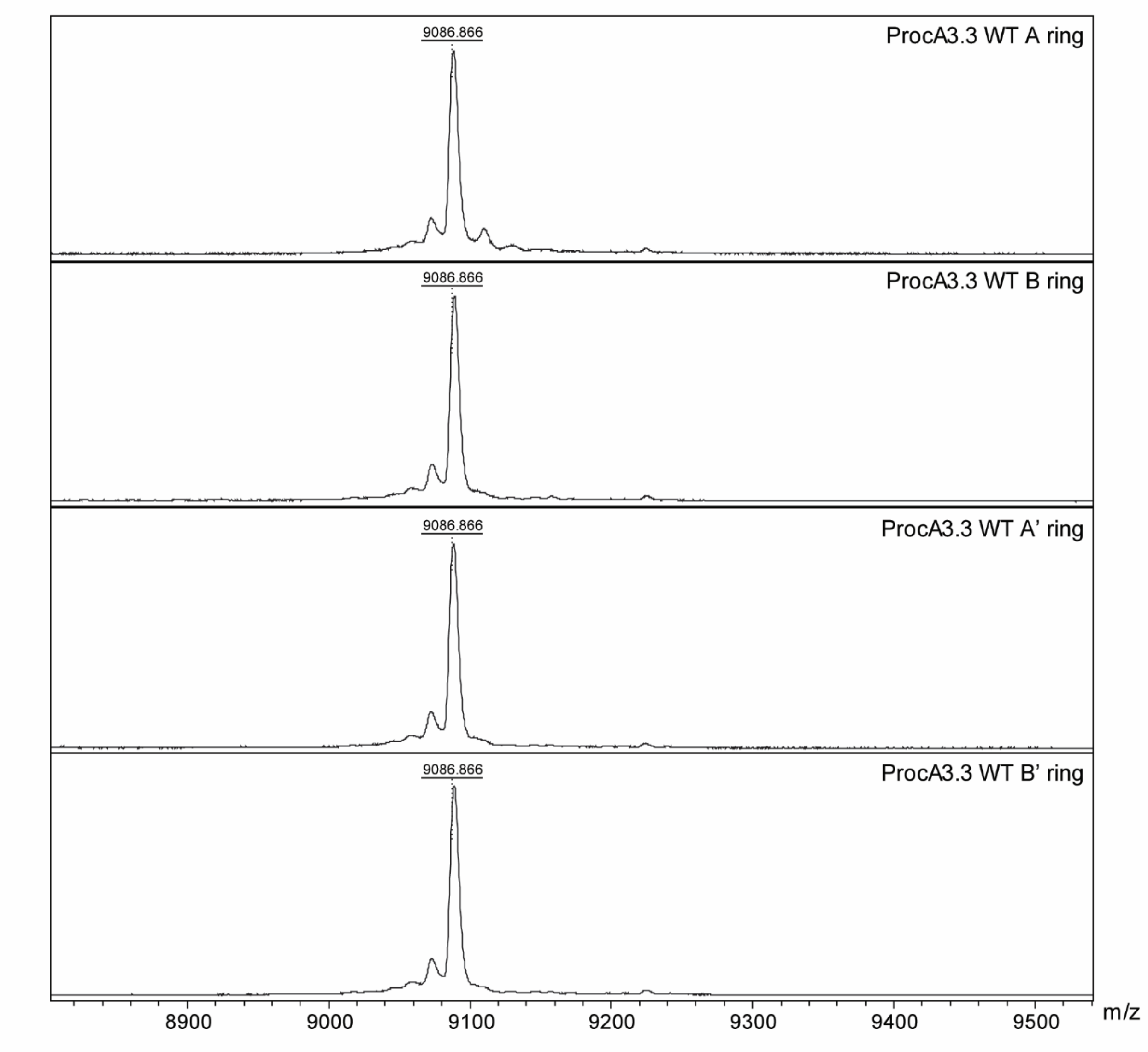

**Figure S16. MALDI-ToF mass spectra for peptides used for the ProcA3.3 WT scaffold isolated ring analysis.** MALDI-ToF mass spectra were taken after peptide purification, thrombin digestion, and HPLC purification of both peptides.

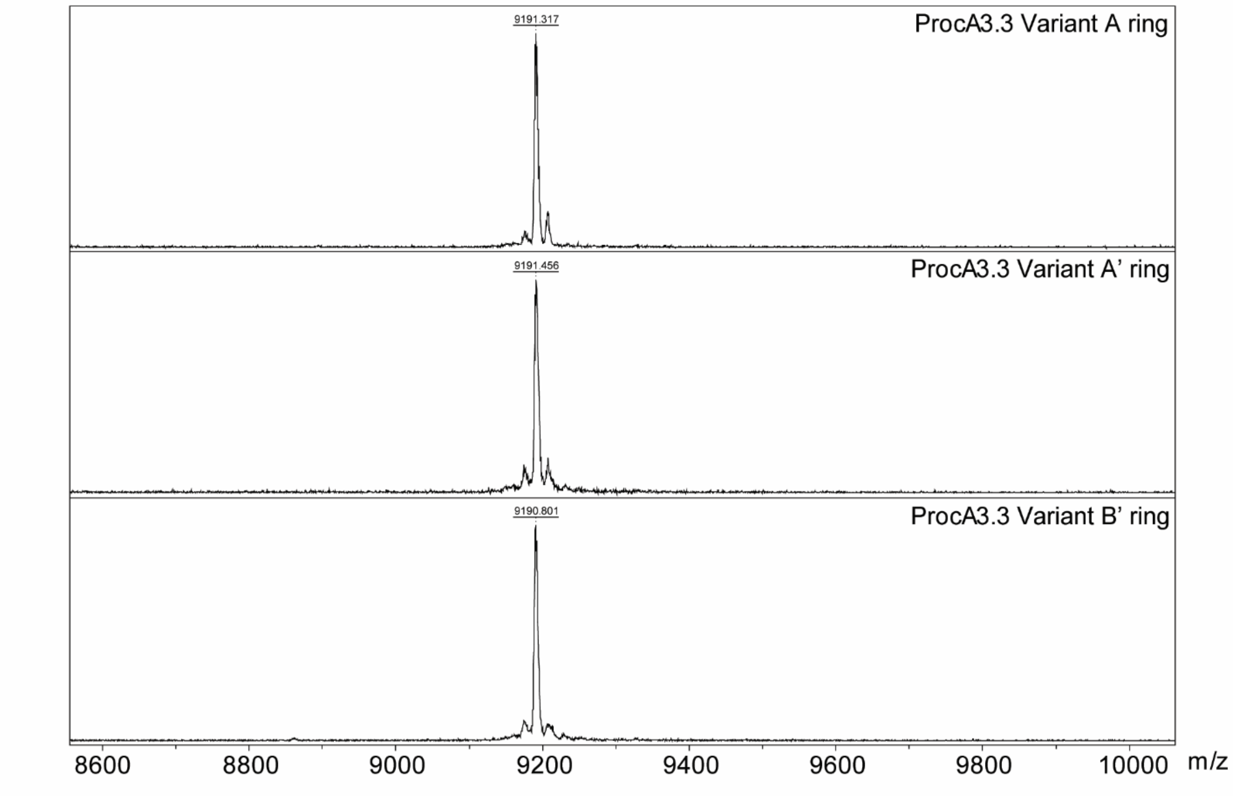

**Figure S17. MALDI-ToF mass spectra for peptides used for the ProcA3.3 variant Z scaffold isolated ring analysis.** MALDI-ToF mass spectra were taken after peptide purification, thrombin digestion, and HPLC purification of both peptides.

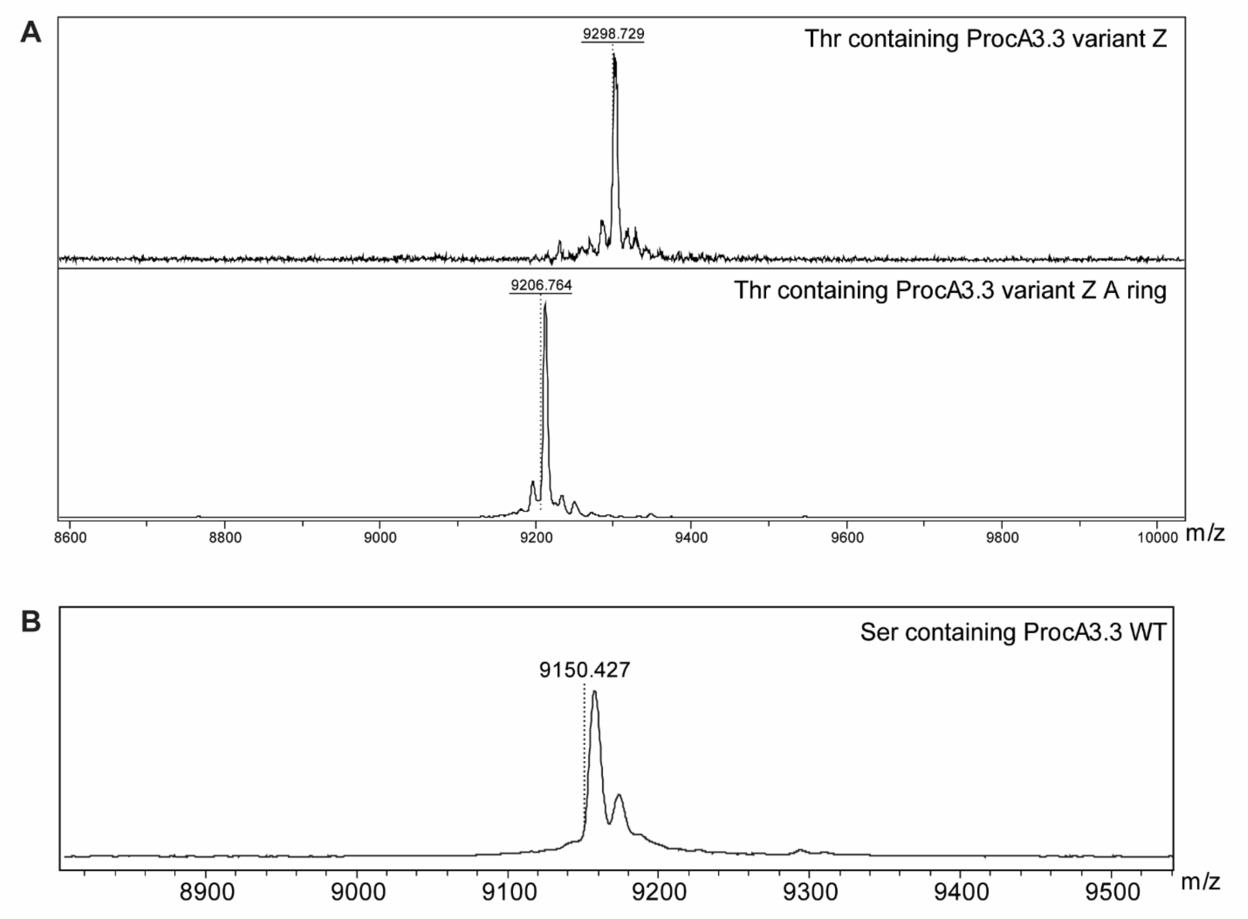

**Figure S18. MALDI-ToF mass spectra for peptides used for Lan vs MeLan analysis. A)** Thr containing ProcA3.3 variant Z peptides. **B)** Ser containing ProcA3.3 WT peptide. MALDI-ToF mass spectra were taken after peptide purification, thrombin digestion, and HPLC purification of both peptides.

**Table S1. Peak assignments for the peptide ions observed in thrombin digested substrates for the indicated ProcM-catalyzed reaction.** Observed masses were determined using deconvoluted spectra.

| Ion Species | Calculated Mass | Observed Mass | Error (ppm) | Reaction conditions, time point |
|---|---|---|---|---|
| **ProcA3.3 WT** | | | | |
| M + 2 NEM + H | 9428.5731 | 9428.5910 | -1.9 | 60 μM, T=20 |
| M + Dhb + 2 NEM + H | 9410.5626 | 9410.5655 | -0.3 | 60 μM, T=20 |
| M + 2 Dhb + 2 NEM + H | 9392.5521 | 9392.5680 | -1.7 | 60 μM, T=20 |
| M + MeLan + NEM + H | 9285.5149 | 9285.5803 | -7.0 | 60 μM, T=20 |
| M + Dhb + MeLan + NEM + H | 9267.5044 | 9267.5683 | -6.9 | 60 μM, T=20 |
| M + 2 Dhb + MeLan + NEM + H | 9249.4939 | 9249.5713 | -8.4 | 60 μM, T=20 |
| M + 2 MeLan + H | 9142.4567 | 9142.5104 | -5.9 | 60 μM, T=20 |
| M + 2 MeLan + Dhb + H | 9124.4462 | 9124.5261 | -8.8 | 60 μM, T=20 |
| **ProcA3.3 variant Z** | | | | |
| M + 2 NEM + H | 9520.59594 | 9520.5427 | 5.6 | 40 μM, T=15 |
| M + Dha + 2 NEM + H | 9502.58544 | 9502.5789 | 0.7 | 40 μM, T=15 |
| M + Lan + NEM + H | 9377.53774 | 9377.4791 | 6.3 | 40 μM, T=15 |
| M + Dha + Lan + NEM + H | 9359.52724 | 9359.4927 | 3.7 | 40 μM, T=15 |
| M + 2 Dha + Lan + NEM + H | 9341.51674 | 9341.4790 | -4.0 | 40 μM, T=15 |
| M + 2 Lan + H | 9234.47954 | 9234.5075 | -3.0 | 40 μM, T=8 |
| M + 2 Lan + Dha + H | 9216.46904 | 9216.4979 | -3.1 | 40 μM, T=8 |
| **ProcA3.3 A ring** | | | | |
| M + Dhb + NEM + H | 9211.5322 | 9211.6028 | -7.7 | 40 μM, T=2 |
| M + NEM + H | 9193.5217 | 9193.5620 | -4.4 | 40 μM, T=2 |
| M + Lan+ H | 9068.474 | 9068.5443 | -7.8 | 40 μM, T=2 |
| **ProcA3.3 variant Z A ring** | | | | |
| M + Dha + NEM + H | 9317.5707 | 9317.6279 | -6.1 | 40 μM, T=2 |
| M + NEM + H | 9299.5602 | 9299.5924 | -3.5 | 40 μM, T=2 |
| M + Lan+ H | 9174.5125 | 9174.5656 | -5.8 | 40 μM, T=2 |
| **ProcA3.3 variant Z MeLan A ring** | | | | |
| M + Dhb + NEM + H | 9331.5863 | 9331.6016 | -1.6 | 40 μM, T=4 |
| M + NEM + H | 9313.5758 | 9313.6098 | -3.6 | 40 μM, T=4 |
| M + Lan+ H | 9188.5281 | 9188.6069 | -8.6 | 40 μM, T=4 |
